# Labile iron overload reprograms microglia and neurons for lipid droplet synthesis in the aging brain

**DOI:** 10.1101/2025.11.03.686315

**Authors:** Karina Cunhae Rocha, Qian Xiang, Chengjia Qian, Lishuan Wang, Wei Yuan, Varsha Beldona, Ying Duan, Luana Veras, Darya Abolmaali, Garam An, Junho Park, Whasun Lim, Xu Chen, Wei Ying

## Abstract

Lipid-droplet (LD) accumulation emerges in microglia and neurons with age. Because neither cell type is specialized for lipid storage, LDs are linked to dysfunction. The upstream drivers of LD formation and their effects on neighboring cells remain unclear. Here, we identify a mitochondrial-iron axis that promotes LD formation in aging microglia via reactive oxygen species (ROS), which secondarily reshapes neuronal iron handling. LD-enriched microglia show reduced mitochondrial mass and increased labile iron, ROS, and lipid peroxidation. Chelating labile iron or scavenging ROS suppresses LD formation. Conditioned media from iron-stressed microglia alter neuronal iron homeostasis, indicating transcellular coupling. In primary neurons, iron overload increases LDs and activates coordinated iron, ROS, and lipogenesis programs, whereas antioxidant treatment attenuates iron-driven LD accumulation. Together, these findings position iron overload as an upstream regulator of ROS-dependent LD biogenesis in microglia and neurons and reveal a microglia-neuron axis that regulates neuronal iron metabolism during aging.

## INTRODUCTION

Diverse microglial states emerge with aging and in neurodegenerative disease, including the lipid-droplet-accumulating microglia (LDAM)^1^. Neurons likewise develop lipid droplets (LDs) with age^2–5^. Because neither microglia nor neurons are specialized for lipid storage, LD accumulation in both cell types is associated with impaired function^1–3^. In microglia, LD accumulation appears to be ROS-dependent^1^, but the full mechanism remains unclear. Moreover, the consequences of microglial LD accumulation for neighboring brain cells, particularly neurons, are still poorly defined.

Microglia originate from embryonic yolk-sac precursors and persist as self-renewing, tissue-resident macrophages in the central nervous system^6–8^. Beyond immune surveillance, microglia are very active cells that maintain brain homeostasis by monitoring neuronal activity, pruning synapses, and phagocytosing myelin, apoptotic neurons, neuronal progenitors, oligodendrocytes, and other cellular debris^9–13^. With aging, microglia undergo functional and metabolic remodeling characterized by reduced process motility, increased pro-inflammatory signaling, elevated reactive oxygen species (ROS), and diminished phagocytic capacity^1,14,15^. Single-cell profiling has further revealed pronounced microglial heterogeneity, including age- and disease-associated states such as disease-associated microglia (DAM), the microglial neurodegenerative phenotype (MGnD), and LDAM^1,16,17^.

Emerging evidence indicates that neurons, in addition to microglia, accumulate LDs during aging and in neurodegenerative disease^2,3^. In both cell types, lipid accumulation correlates with elevated ROS^1,18,19^. Excess intracellular iron is a well-established trigger of ROS production^20–23^. Iron is the most abundant transition metal in the brain and is essential for neuronal and glial function^24–26^. Mouse and human studies consistently report increased brain iron with aging and in multiple neurodegenerative conditions, with some studies showing iron overload as a trigger for ROS production and associated with Alzheimer’s disease (AD) and Parkinson’s disease (PD)^20,27–35^ . In vitro studies indicate that microglia are particularly efficient at iron sequestration, with microglial iron stores approximately threefold higher than those of neurons^36^. Post-mortem analyses in PD, multiple sclerosis, and amyotrophic lateral sclerosis further support microglia as key iron-sequestering cells in affected regions^37–39^. While the cause of iron accumulation in the aging brain is unclear, evidence indicates that expression of ferroportin (*Fpn*/*Slc40a1*), which exports iron, and ferritin heavy chain 1 (*Fth1*) and ferritin light chain 1 (*Ftl1*), which store iron, changes with aging and with neurodegenerative disorders^29,32,40,41^.

It remains unknown whether iron accumulation can act upstream of ROS to promote lipid-droplet formation in microglia and neurons and whether microglial iron handling calibrates neuronal iron status and lipid remodeling. We addressed this in mouse hippocampus using tamoxifen-inducible, microglia-specific knockouts of *Fth1* and *Fpn*, mechanistic assays in BV2 microglia with iron and LPS challenges plus antioxidant or iron-chelation rescue, and primary neuron models exposed to oleic acid and ferric ammonium citrate (FAC), supported by bulk RNA sequencing (RNA-seq), single-cell RNA-seq (scRNA-seq), single-nucleus RNA-seq (snRNA-seq), and conditioned-media experiments. Our results define microglia-neuron iron axis that links iron availability to ROS and lipid-droplet programs in aging.

## RESULTS

### Lipid droplet-enriched microglia are Fe^2+^ overloaded

LD accumulation in microglia has been well described, but the mechanisms leading to that accumulation remain unclear. To investigate this, we analyzed bulk RNA-seq from mouse hippocampus (GSE208386; young 3 months vs aging 16 months, both sexes) and single-nucleus RNA-seq from human hippocampus (GSE18553 and GSE199243 young 18-39 years vs aging 60-95 years, both sexes) and performed pathway enrichment analyses to identify age-associated differences^42,43^. Interestingly, alongside the expected strong activation of lipid-metabolism pathways, microglia in the aging brain were characterized by enrichment of pathways related to mitochondrial dysfunction (**Figure S1A**). To more directly explore pathways associated with lipid accumulation, we reanalyzed a bulk RNA-seq dataset (GSE139542) to assess pathway enrichment in microglia with high versus low levels of LD isolated from 18-month-old male mouse hippocampi^1^. In the aging hippocampus, pathways associated with mitochondrial dysfunction, response to oxidative stress, and lipogenesis were upregulated in microglia enriched with LD (**Figure 1A**). Based on in silico analyses implicating mitochondrial dysfunction in aging and LD-enriched microglia, we assessed mitochondrial mass in microglia from young and aged mice by flow cytometry using MitoTracker dye. Consistent with the in silico findings, mitochondrial mass was markedly reduced in microglia during aging (**Figure 1B**). In parallel, neutral lipids were labeled with LipidSpot 488 to define LD status, revealing that LD⁺ microglia exhibited lower mitochondrial mass than LD⁻ microglia (**Figure 1C**). Consistent with the mitochondrial mass changes, ferritin, a marker of intracellular iron content, was one of the upregulated genes in aging microglia in both bulk RNA-seq (GSE208386) and single-cell RNA-seq (scRNA-seq, GSE161340) analyses of the mouse hippocampus (**Figures 1D&S1B**)^42,44^. We next assessed the impact of aging on iron phenotypes in microglia. Given that labile iron supply is essential for mitochondrial fitness, we quantified labile iron content in LD⁻ and LD⁺ microglia across ages using a fluorescent probe that selectively reacts with Fe²⁺ and converts it into a far-red fluorescent substance^45^. By 36 weeks of age, approximately 50% of microglia were LD- enriched, a pattern that was maintained at 44 and 94 weeks of age (**Figure S1C**). LD⁺ microglia consistently displayed significantly higher labile iron levels than LD⁻ microglia across all age groups (**Figures 1E-G**). Furthermore, aged LD⁺ microglia showed marked iron overload compared with their young counterparts (**Figures 1E-G**). In bulk RNA-seq, the LD-high microglia showed coordinated induction of *Trf*, *Ftl*, and *Slc40a1*, together with *Rab7*, *Atg3*, and *Lamp1* (late endosome/lysosome and autophagy machinery) (**Figure S1D**). This transcriptional signature indicates enhanced transferrin-dependent endosomal iron trafficking and activation of ferritinophagy, which involves the release of Fe^2+^ from ferritin. As Fe²⁺ accumulation can induce ferroptosis, we evaluated whether aging microglia show ferroptotic features by measuring MDA, an end product of lipid peroxidation of cell membranes, by flow cytometry^46^. As shown in **Figure 1H**, aging microglia showed higher levels of MDA. When comparing LD⁺ versus LD⁻ in the aging brain, as expected, LD⁺ microglia had approximately three-fold higher MDA than LD⁻ microglia (**Figure 1I**). Bulk RNA-seq analysis showed no difference in Gpx4 and Slc7a1 expression between LD⁺ and LD⁻ microglia, suggesting that lipid and iron accumulation occur without a compensatory anti-ferroptotic response (**Figure S1E**). Additionally, scRNA-seq analysis indicates low or barely detectable Gpx4 and Slc7a11 expression in mouse hippocampal microglia, suggesting limited anti-ferroptotic capacity (**Figure S1F**). Corroborating these observations, per-sample pathway scores (mean expression of pathway genes) were higher for ferroptosis and oxidative stress in LD⁺ than in LD⁻ microglia (**Figure 1J**). Together, these findings identify a mitochondria-iron axis in aging LD-enriched microglia characterized by low mitochondrial mass, ferritin upregulation with labile iron excess, and increased lipid peroxidation in the setting of limited anti-ferroptotic programs.

**Figure 1.**
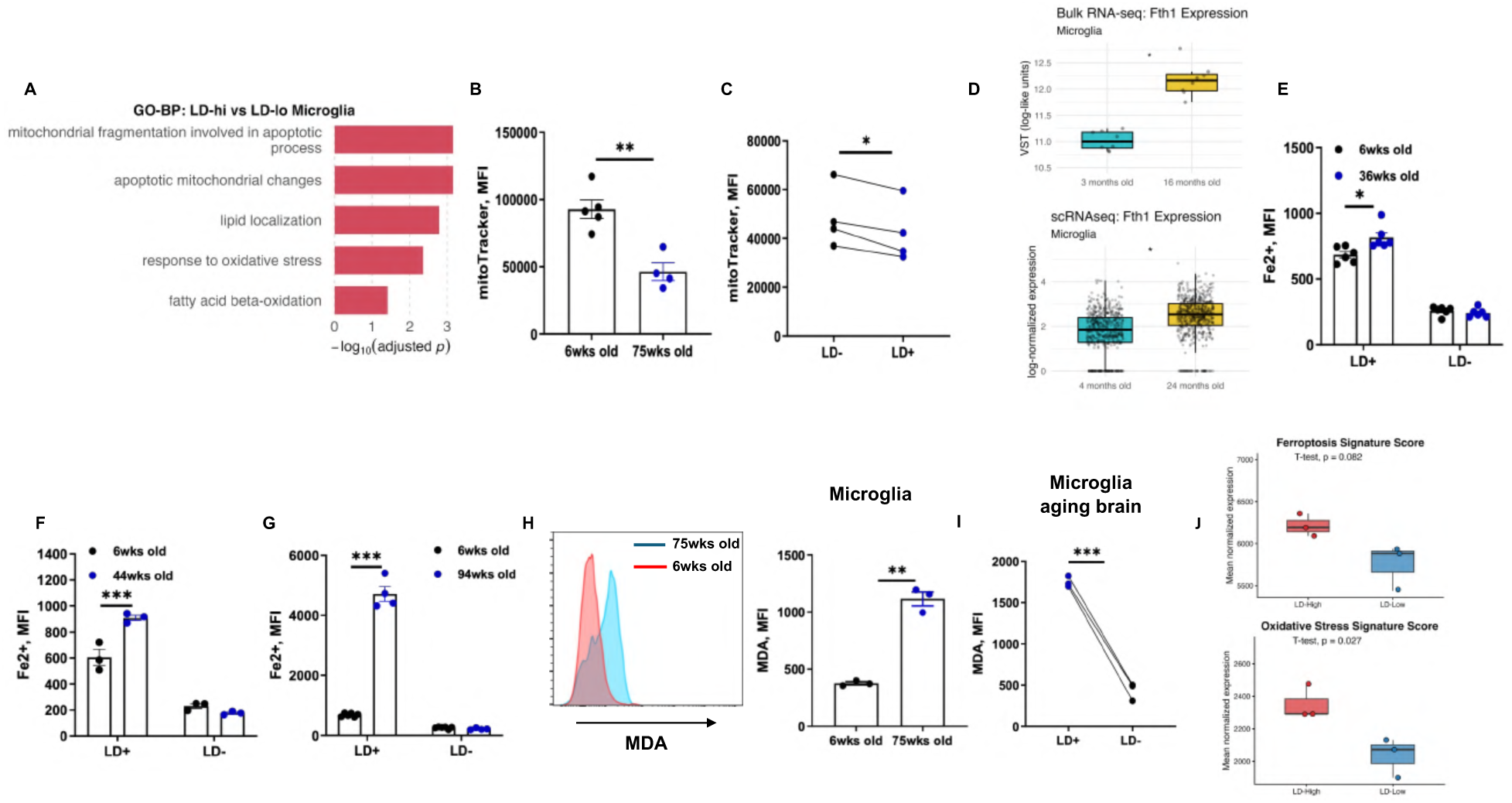
Lipid-droplet-enriched microglia accumulate Fe²⁺ and exhibit oxidative stress in aging hippocampus. **(A)** Gene Ontology Biological Process (GO-BP) enrichment from bulk RNA-seq of LD-high vs LD-low microglia isolated from 18-month mouse hippocampus. **(B)** Flow-cytometric mitochondrial mass in hippocampal microglia from 6- vs 75-week-old mice, quantified as MitoTracker mean fluorescence intensity (MFI). **(C)** MitoTracker MFI in LD⁺ vs LD⁻ microglia within aged mice. **(D)** Ferritin heavy chain (*Fth1*) expression in microglia from mouse bulk RNA-seq and scRNA-seq comparing young vs aging; plots show normalized counts per dataset.**(E-G)** Labile Fe²⁺ content in microglia at 6, 36, 44, and 94 weeks of age, stratified by LD status (LD⁺/LD⁻), measured by an Fe²⁺-reactive fluorescent probe via flow cytometry. **(H)** Lipid peroxidation in microglia assessed by malondialdehyde (MDA) fluorescence. **(I)** MDA MFI in LD⁺ vs LD⁻ microglia within aging mice. **(J)** Per-sample module scores for ferroptosis and oxidative-stress gene sets in LD⁺ vs LD⁻ microglia. Points represent biological replicates; bars, mean ± s.e.m. Two-group contrasts used two-tailed unpaired Student’s t-tests. Matched LD⁻/LD⁺ comparisons used paired t-tests; multi-factor effects (age x LD status) used two-way ANOVA with Tukey post-hoc. Bulk RNA-seq DE was computed with DESeq2 (Wald test) and Benjamini-Hochberg FDR, while scRNA-seq gene tests used Wilcoxon rank-sum with FDR control. Exact P values are shown when available; otherwise *P ≤ 0.05, **P ≤ 0.01, ***P ≤ 0.001.

### Labile iron overload induces aging-related microglia abnormalities through ROS accumulation

Previous studies have shown that 40-week-old PS19 mice, a mouse model that induces the tau pathology seen in AD, exhibit a marked reduction in microglial number alongside increased microglial lipid accumulation^19^. Because this mirrors what we observed in the aging phenotype, we asked whether PS19 microglia also exhibit changes in iron metabolism as seen in aging mice microglia. As shown in **Figure S2A**, LD⁺ microglia isolated from 40-week-old PS19 mice displayed elevated labile Fe²⁺. Notably, we previously observed Fe²⁺ overload in LD+ microglia of WT mice by 36 weeks of age (**Figures 1E-G** **& S1C**). Therefore, we next assessed whether microglial LD accumulation and iron overload occur independently of age in the PS19 model.

For that, we compared microglia phenotypes between PS19 mice and their WT littermates at 8 weeks of age. Both groups have comparable microglia numbers and Fe^2+^ levels (**Figures 2A&S2B**), indicating that early tau pathology has minimal impact on the microglial iron phenotype. Previous studies have demonstrated that activation of lipopolysaccharide (LPS)- mediated pathways contribute to aging-related microglial abnormalities^1^. Consistent with these reports, 24-hour LPS treatment induced LD accumulation in the BV2 microglial cell line (**Figure S2C**). Similar to microglia from the aging brain, lipid accumulation in BV2 cells is accompanied by changes in iron metabolism. LD⁺ BV2 cells (LPS-treated) contained higher Fe²⁺ levels and higher expression of *Fth1* than LD⁻ BV2 cells (**Figures 2B and 2C**). In contrast, LPS treatment downregulated the iron importer *Tfrc1* and the iron exporter *Fpn*, suggesting compensatory responses to rebalance iron content (**Figure 2C**). In addition, LD⁺ BV2 cells had more mitochondrial Fe²⁺ than LD⁻ BV2 cells (**Figure 2D**). To validate the impact of LPS on microglial iron phenotypes in vivo, a group of 8-week-old WT mice were injected intraperitoneally with LPS (1 mg/kg BW) for 3 days. As shown in **Figures 2E&2F**, microglia exhibited increased Fe²⁺ and ROS in response to LPS stimulation. In addition, microglia isolated from LPS-treated mice expressed higher *Fth1* than those from control mice (**Figure S2D**). We also observed that LPS injection resulted in greater MDA levels, concomitant with a reduction in microglia population (**Figures 2G&S2E**). To assess the importance of Fe²⁺ supply for LD synthesis in microglia, BV2 cells were co-treated with LPS and the labile iron chelator 2,2′-bipyridine (Bipy)^47,48^. As shown in **Figure 2H**, Bipy significantly blocked LPS-induced LD synthesis in BV2 cells. Additionally, expression of the acetyl-CoA carboxylase (*Acc*) gene, which catalyzes the rate-limiting step in fatty-acid synthesis, was upregulated after LPS stimulation, potentially driving the observed lipid accumulation (**Figure S2F**). Bipy normalized *Acc* expression to basal levels, further supporting the role of labile iron in lipid regulation (**Figure S2F**). Conversely, iron overload induced by ferric ammonium citrate (FAC; 100 µM) increased *Fth1* expression (**Figure S2G**), and LDs were readily detected after 24 hours in BODIPY-stained BV2 cells (**Figure 2I**).

**Figure 2.**
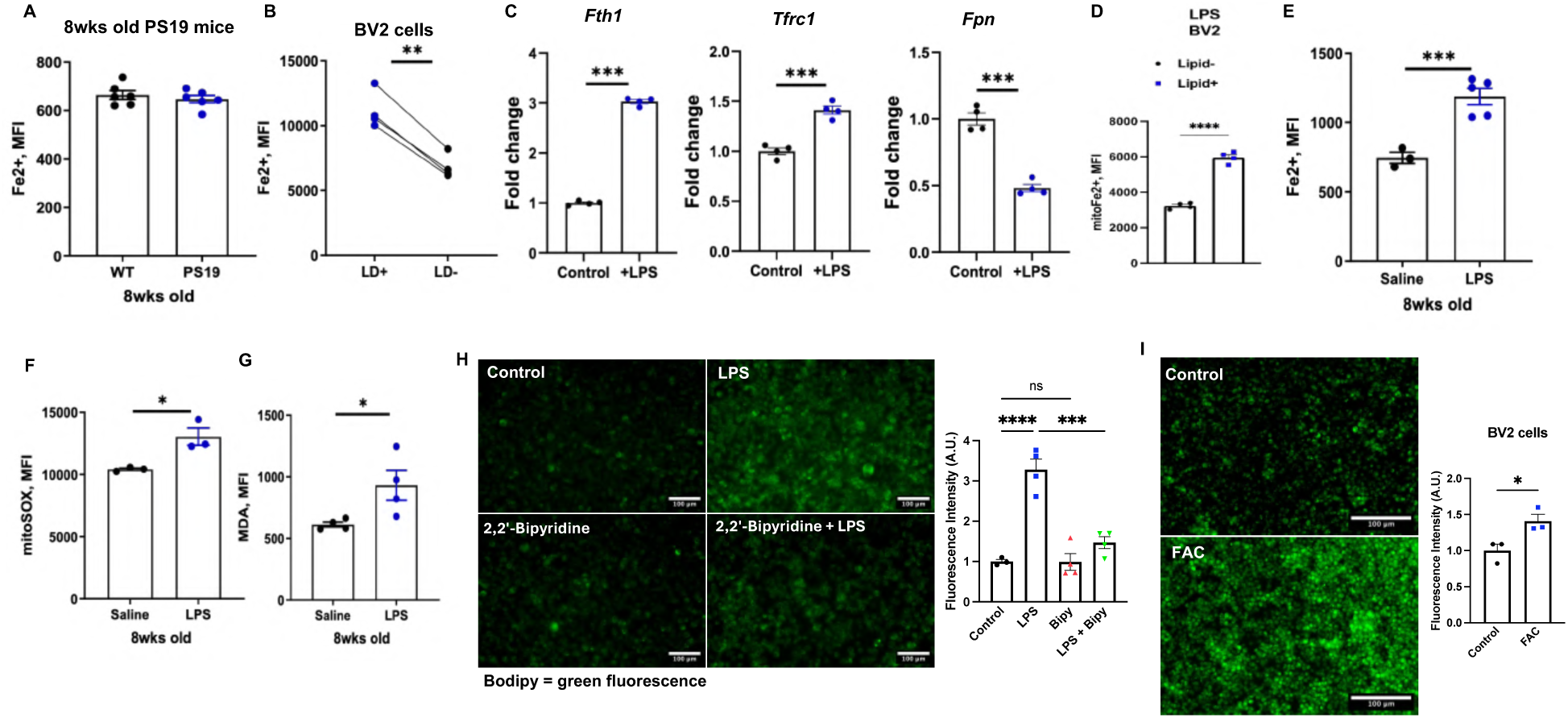
Labile iron and LPS drive lipid-droplet formation and oxidative stress in microglia. **(A)** Microglial labile Fe²⁺ mean fluorescence intensity (MFI) in 8-week-old WT vs PS19 mice by flow cytometry. **(B)** BV2 labile Fe²⁺ MFI in LD⁺ vs LD⁻ populations after LPS (5 µg) treatment. **(C)** Expression of *Fth1*, *Tfrc1*, and *Slc40a1* (*Fpn*) in BV2 cells treated with LPS (5 µg) or control (saline) for 24 h. **(D)** Mitochondrial Fe²⁺ MFI in LD⁺ vs LD⁻ BV2 cells after LPS treatment. **(E- G)** Microglial **(E)** labile Fe²⁺, **(F)** mitochondrial ROS, and **(G)** lipid peroxidation (MDA) MFI in 8-week-old mice after saline or LPS injections (1 mg/kg once daily for 3 days). **(H)** BODIPY fluorescence images and quantification (arbitrary units, A.U.) of BV2 cells after 24 h treatment with control (saline), LPS (5µg), 2,2′-bipyridine (Bipy, 50 µM), and LPS+Bipy. **(I)** BODIPY images and quantification (A.U.) of BV2 cells after 24 h treatment with control (saline) or ferric ammonium citrate (FAC, 100 µM). Points represent biological replicates; bars, mean ± s.e.m. Two- group comparisons used two-tailed unpaired Student’s t-test; multi-group comparisons used one- way ANOVA with Tukey’s post-hoc test. *P ≤ 0.05, **P ≤ 0.01, ***P ≤ 0.001. scale bar, 100 µm.

### Labile iron overload couples to ROS and lipid-droplet (LD) formation in aging microglia

It is well studied that ROS accumulation is a key driver of lipogenesis^1,18^. Here, concomitant with high Fe²⁺ content and changes in mitochondrial function in LD⁺ microglia, ROS production was higher in LD⁺ than LD⁻ cells at both 6 and 75 weeks, with a further increase in LD⁺ microglia at 75 weeks (**Figure 3A**). In BV2 cells, 24 hours of LPS stimulation increased mitochondrial ROS preferentially in LD⁺ cells relative to LD⁻/control cells (**Figure 3B**). Total cellular ROS was also increased by LPS in BV2 cells, and iron chelation with Bipy (50 µM) during LPS stimulation prevented this LPS-induced increase in cellular ROS (**Figure 3C**). In addition, removal of ROS by N-acetylcysteine treatment (NAC; 1mM/well) blunted the ability of LPS to induce LD synthesis in BV2 cells (**Figure S3A**). Similarly, NAC treatment significantly blocked LD synthesis in response to FAC (**Figure 3D**), implicating ROS as a key mediator linking iron overload to microglial lipogenesis. We next asked whether analogous programs exist in microglia from aging (24 months old) mouse hippocampus profiled by scRNA-seq (GSE161340)^44^. Re-analysis and integration of public hippocampal scRNA-seq (LogNormalize/RPCA) identified microglia by canonical markers (*P2ry12*, *Tmem119*, *Cx3cr1*, and *Aif1*) occupying discrete UMAP neighborhoods across ages (**Figure S3B**). Within aged microglia, stratification by *Fth1* expression into quartiles revealed an enrichment for lipid- metabolism and oxidative stress response pathways in Fth1-High (Q4) cells, compared with Fth1-Low (Q1) (**Figures 3E&S3C**). Together, these findings link iron handling to a coordinated lipid-anabolic/oxidative-stress state in microglia from aging mice, mirroring the iron/ROS- dependent LD accumulation phenotype observed in BV2 cells. Building on the in vitro and in silico findings, we next tested the phenotype in vivo, using young animals. We generated a tamoxifen-inducible, microglia-specific *Fth1* knockout mouse (Tmem119^ERT2Cre^/Fth1^flox/flox^; **Figure S3D**) aiming to promote iron accumulation in microglia in young mice. As expected, loss of *Fth1*, critical for iron storage via sequestration of labile iron, resulted in elevated microglial Fe²⁺ levels compared with WT littermates (**Figure 3F**). Consistent with this, mitochondrial Fe²⁺ was higher in Fth1KO microglia than in WT, accompanied by increased mitochondrial ROS production (**Figure 3G&H**). Additionally, *Fth1* deficiency increased the proportion of LD^+^ microglia (**Figure 3I**). Together, these data indicate that labile iron accumulation drives ROS production alongside lipid accumulation in aging microglia.

**Figure 3.**
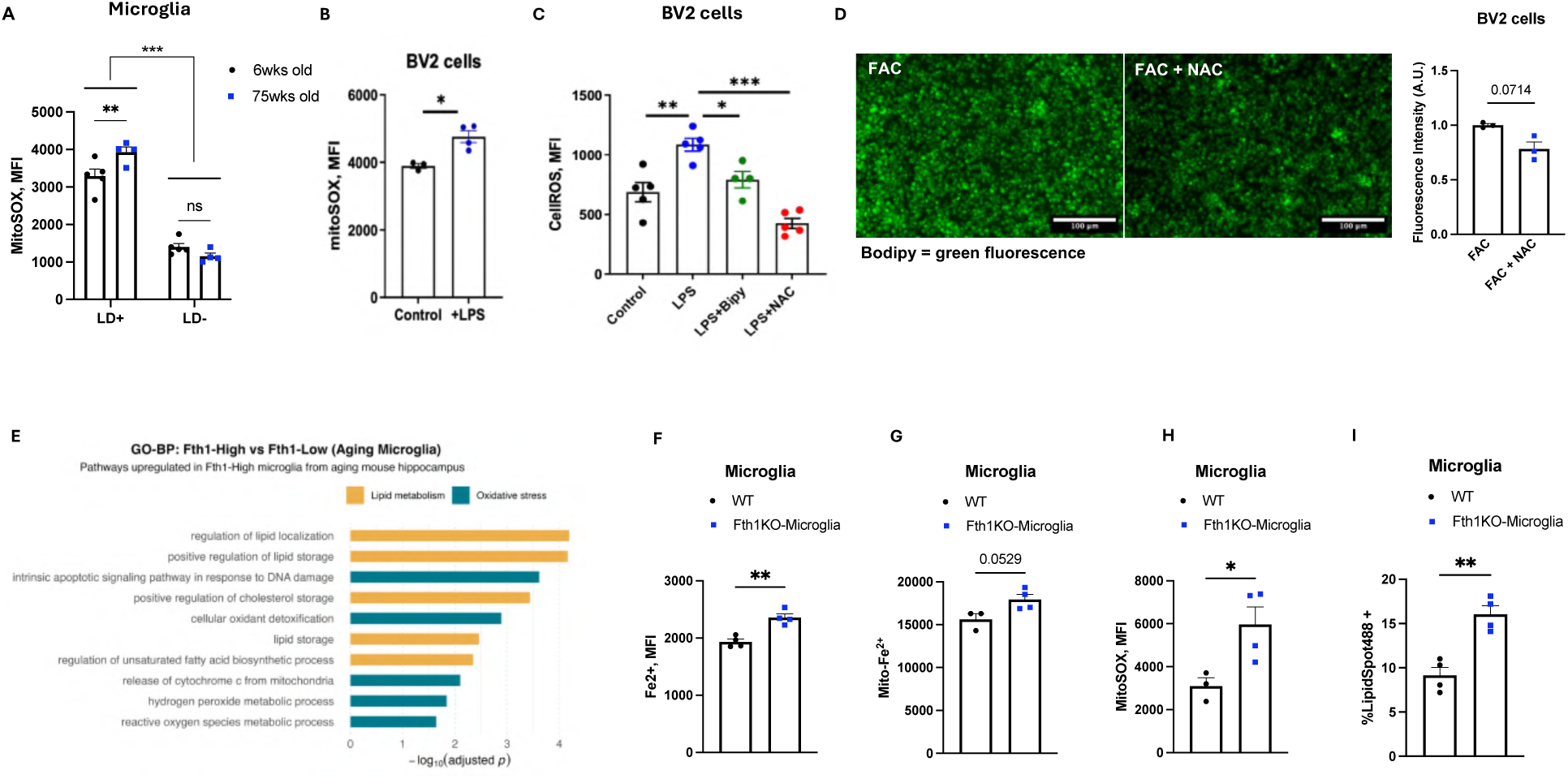
Labile iron overload couples ROS to lipid-droplet formation in aging microglia. **(A)** Mitochondrial ROS in brain microglia from 6- and 75-week-old mice, shown as MitoSOX mean fluorescence intensity (MFI) and split by LD status. **(B)** BV2 mitochondrial ROS after 24 h treatment with control (saline) or LPS (5 µg). **(C)** BV2 total cellular ROS after 24 h treatment with control, LPS (5 µg), LPS+2,2′-bipyridine (Bipy, 50 µM), or LPS+N-acetylcysteine (NAC, 1mM). **(D)** Representative BODIPY images and fluorescence quantification (arbitrary units, A.U.) of BV2 cells treated 24 h with ferric ammonium citrate (FAC) with or without NAC. **(E)** Gene Ontology Biological Process (GO-BP) enrichment from re-analysis of 24-month mouse hippocampal microglia (scRNA-seq): Fth1-High (quartile 4) vs Fth1-Low (quartile 1). **(F)** Labile Fe²⁺, **(G)** mitochondrial Fe²⁺, **(H)** mitochondrial ROS, and **(I)** percentage of LipidSpot⁺ of microglia isolated from tamoxifen-inducible, microglia-specific *Fth1* knockout vs WT mice. Points represent biological replicates; bars, mean ± s.e.m. Two-group comparisons (**B**, **D**, **F–I**) used two-tailed unpaired Student’s t-test. Multi-group comparisons (**C**) used one-way ANOVA with Tukey’s post hoc test. Panel (**A**) used two-way ANOVA (age × LD status) with Tukey’s post hoc test. *P ≤ 0.05, **P ≤ 0.01, ***P ≤ 0.001. Scale bars, 100 µm.

### Aging reprograms neurons and increases neuron-microglia interactions

Motivated by iron dysregulation in aging microglia, we asked whether neurons exhibit similar alterations. We processed publicly available mouse hippocampus (dentate gyrus) scRNA-seq datasets from young (9 weeks old) and aging (16-21 months old) mice and resolved major cell populations using SCT-normalized PCA/UMAP (**Figure S4A**, GSE233363)^49^. Canonical markers confirmed identities of neurons (*Snap25*, *Rbfox3*, *Tubb3*, *Map2*) and microglia (*C1qa*, *Tmem119*, *P2ry12*, *Cx3cr1*), which segregated into distinct UMAP clusters (**Figures S4B-D**). Age-split embeddings showed that these identities remain transcriptionally separable in both young and aged samples, with age-dependent shifts in their distributions (**Figure 4A**). Genes upregulated in aging neurons were enriched for GO Biological Process terms related to metal-ion/iron transport, oxidative-stress/ROS responses, and lipid catabolism, mirroring the iron– lipid–ROS signature seen in aging microglia (**Figure 4B**). These observations prompted us to test the interaction between neuron-microglia. To test whether neuron-microglia interactions and spatial organization change with age, we analyzed published Visium spatial transcriptomics from young and aging mouse hippocampus (GSE233363)^49^. Spots were embedded with UMAP, clustered using a graph-based approach, and annotated as neuronal or microglial based on canonical markers (*Snap25*, *C1qa*) (**Figure S4E**). We then used the neuron and microglia spot populations to compute neuron-microglia distances from tissue coordinates (pixels converted to µm), enabling age-stratified proximity analyses. As shown in **Figure 4C**, neuron-microglia distances were significantly reduced in the aged hippocampus. We then asked whether proximity relates to neuronal transcriptional remodeling. Neurons in the closest quartile to microglia show GO-BP enrichment for metal-ion transport, lipid remodeling (DAG metabolism, lipid phosphorylation, glycerolipid/glycerophospholipid biosynthesis), and stress/ROS signaling relative to the farthest quartile (**Figure 4D**). As a complementary approach, we applied CellChat to the scRNA-seq dataset (GSE233363) restricted to neurons and microglia ^49^. Separate CellChat models for young and aging and compared revealed an increase in microglia-neuron signaling from 6 to 11 significant ligand-receptor pairs, with a shift toward cell-cell contact and ECM-receptor classes alongside secreted cues (**Figure 4E**). Pair-level inspection revealed aging-specific gains in CADM1-CADM1 (adhesion), PTPRC-CD22 and VSIR-IGSF11 (immune/checkpoint tone), and SEMA6A-PLXNA2/4 (axon-guidance), as well as stronger shared programs such as PDGFB-PDGFRB, HSPG2/LAMC1-DAG1, and CDH2-CDH2 (**Figure 4F**). Collectively, the proximity-linked pathway enrichments and the aging-associated increase in contact/ECM ligand-receptor signaling support a model in which microglia-neuron interactions intensify with age and align with neuronal programs related to metal-ion/iron transport, lipid remodeling, and oxidative stress in mouse. We next asked whether these pathways are similarly enriched in aging human neurons. Using two publicly available human hippocampal datasets (GSE185553 and GSE199243) using SCT-normalized PCA/UMAP clustering (**Figure S4E**) ^43^.

**Figure 4.**
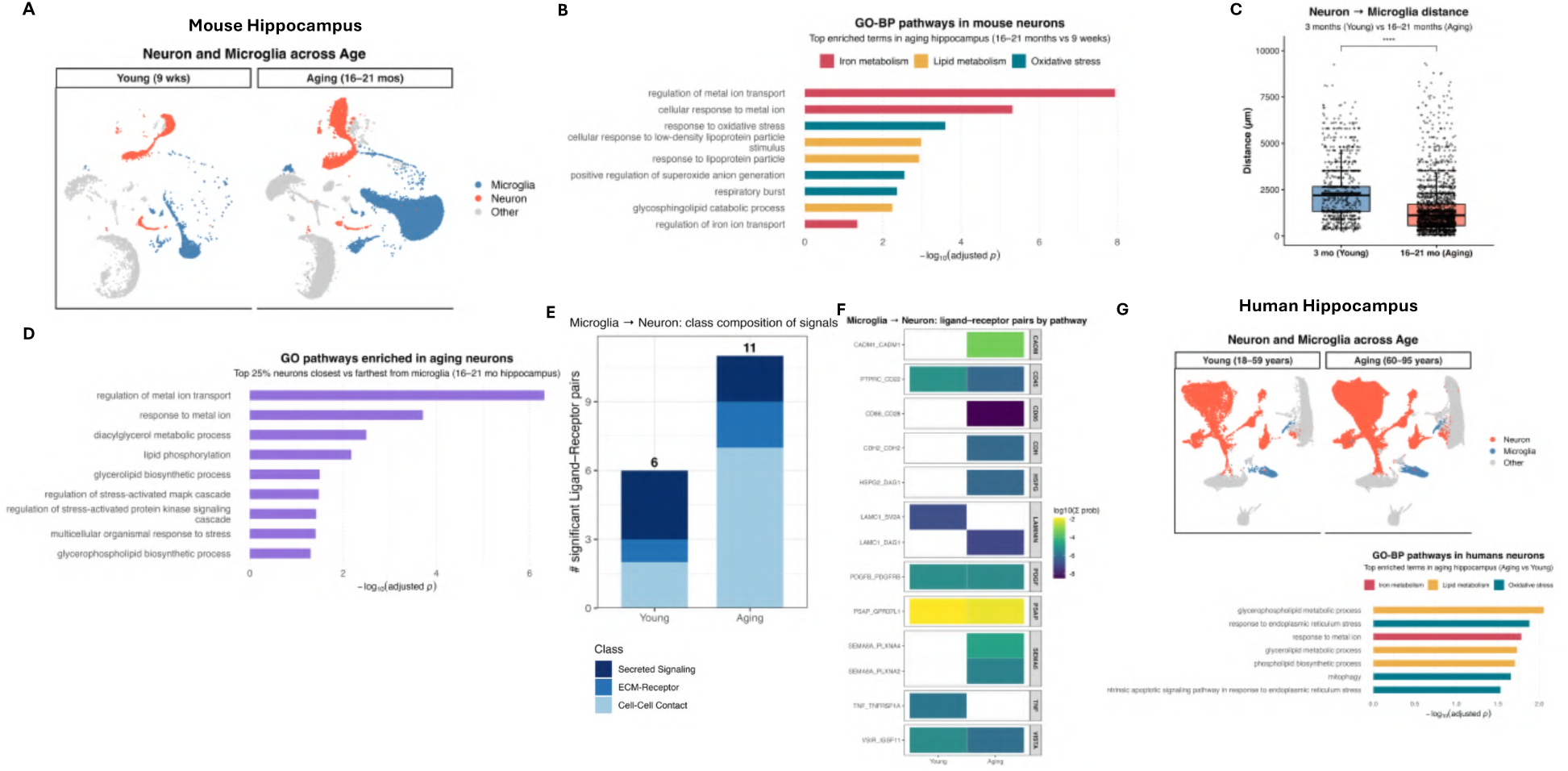
Aging reprograms neurons and intensifies neuron-microglia interactions. **(A)** Mouse dentate gyrus scRNA-seq: UMAPs split by age (Young, 9 weeks old; Aging, 16-21 months old) showing Neuron, Microglia, and Other populations defined by canonical markers. **(B)** GO- Biological Process enrichment for genes upregulated in aging mouse neurons (Aging vs Young), grouped into metal-ion/iron handling, lipid metabolism, and oxidative-stress programs. **(C)** Visium spatial transcriptomics: neuron-to-microglia nearest-neighbor distances (µm) computed from tissue coordinates and compared between Young and Aging hippocampus. **(D)** GO-BP enrichment in mouse neurons stratified by proximity to microglia (top-quartile Near vs top-quartile Far), highlighting metal-ion transport, lipid remodeling, and stress/ROS pathways. **(E)** CellChat analysis restricted to neurons and microglia: number of significant ligand-receptor interactions by class in Young vs Aging. **(F)** Selected ligand-receptor pairs illustrating aging-biased interactions and conserved signals. **(G)** Human hippocampus snRNA-seq integration: UMAPs split by age (Young, 18-39 years old; Aging, 60-95 years old) showing Neuron and Microglia subsets. **(H)** GO-BP enrichment for genes upregulated in aging human neurons (Aging vs Young), highlighting metal-ion/iron transport, lipid metabolic processes, and ER-stress/ROS pathways. For bar/box plots, points denote samples/cells/spots as indicated; bars show mean (or the plotted enrichment statistic). Distance and group comparisons used Wilcoxon tests; GO terms were FDR-corrected (Benjamini-Hochberg).

The UMAP embedding resolved transcriptionally distinct clusters across the integrated samples spanning young (18-39 years old) and aging (60-95 years old) adults and focused on neurons and microglia (**Figure 4G**). UMAP visualization and canonical markers confirmed robust identification of neuronal and microglial populations across ages (**Figures S4G&S4H**).

Differential expression in neurons revealed GO-BP enrichment for metal-ion transport, lipid remodeling pathways (glycerophospholipid/glycerolipid metabolism, phospholipid biosynthesis), and stress/ROS programs (ER-stress responses and related apoptotic signaling) (**Figure 4H**). Together, mouse spatial/scRNA-seq and human snRNA-seq analyses support a model in which, similar to what was observed in microglia, aging drives neuronal reprogramming toward iron/metal-ion transport, lipid metabolic processes, and oxidative-stress responses.

### Iron-stress microglia impact neuronal iron handling

We are not the first one to observe iron enrichment in the brain as an aging characteristic. In peripheral tissues, tissue-resident macrophages (e.g., Kupffer cells) play a critical role in governing microenvironmental iron homeostasis^50^, thus we investigated whether microglia might play a similar role in the brain. snRNA-seq analysis (GSE161340) indicates that microglia have higher iron levels than neurons in both young and aged mice, as evidenced by higher ferritin abundance at the single-cell level and in pseudobulk analysis (**Figures 5A&S5A**)^44^. With aging, ferritin expression increased in both microglia and neurons (**Figures 1D&5B**), and cell-cell communication analysis suggested enhanced microglia-neuron crosstalk (**Figures 4E&4F**). We therefore tested whether microglia regulate neuronal iron levels. To reduce microglial iron export, we generated a tamoxifen-inducible, microglia-specific ferroportin KO (*Fpn*/*Slc40a1*; **Figure S5B**). Notably, *Fpn* loss produced minimal changes in microglial numbers and labile Fe²⁺ within microglia (**Figures 5C&5D**). In contrast, hippocampal FTH1 abundance was reduced in FpnKO mice, which could suggest lower total iron content in cells other than microglia (**Figure 5E**). To test whether microglia influence neuronal iron, we exposed primary neurons to microglia conditioned media (CM). Neuron were isolated from postnatal day (PD) 0-2 pups, maintained for 12-15 days in vitro (**Figure S5C**), and then treated with CM collected from microglia-specific WT or FpnKO mice. Neuronal ferritin heavy chain (*Fth1*) was unchanged, whereas ferritin light chain (*Ftl*) tended to decrease after exposure to FpnKO CM (**Figures 5F&5G**). In a complementary experiment, neurons treated with CM from microglia-specific Fth1KO mice showed higher neuronal *Fth1* expression than those treated with WT CM (**Figure 5H**). Additionally, when primary neurons were treated with CM from LD⁻ versus LD⁺ microglia, *Fth1* levels were higher in neurons receiving LD⁺ microglia CM (**Figure 5I**). Together, these findings indicate that aging-associated microglial states not only remodel microglial iron metabolism but also influence neuronal iron handling.

**Figure 5.**
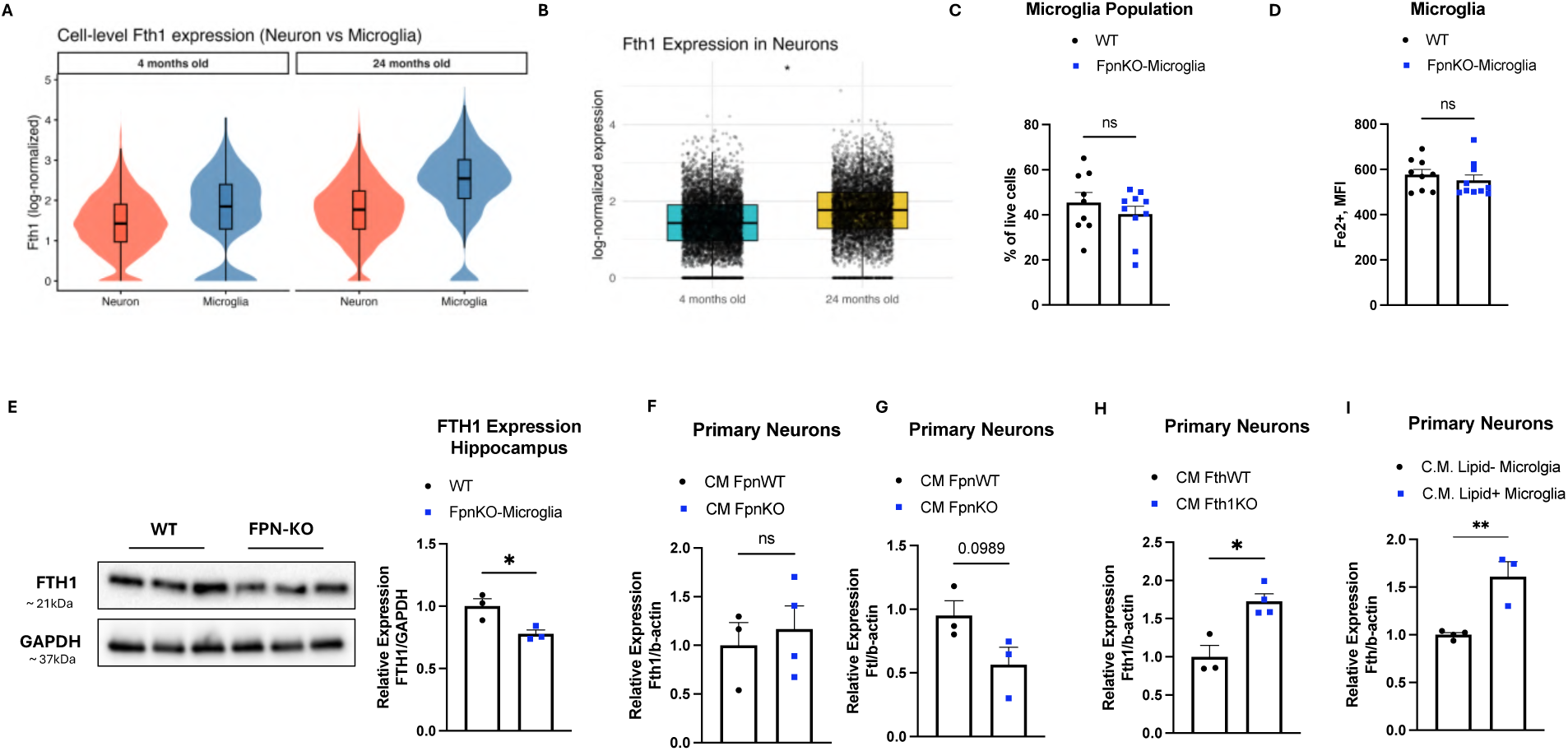
Microglial iron levels influence neuronal iron homeostasis. **(A)** Single-cell *Fth1* expression in neurons vs microglia at 4 and 24 months in mouse hippocampus (snRNA-seq); violin/boxplot overlays show log-normalized counts per cell. **(B)** Neuronal *Fth1* expression (snRNA-seq) comparing 4 vs 24 months. **(C-D)** Tamoxifen-inducible, microglia-specific ferroportin knockout (FpnKO-Microglia) vs WT littermates. **(C)** percentage of microglia among live brain cells measured by flow cytometry; **(D)** microglial labile Fe²⁺ mean fluorescence intensity (MFI). **(E)** Whole-hippocampus immunoblot of ferritin heavy chain (FTH1) with band-intensity quantification in WT vs FpnKO mice. **(F-G)** Expression of **(F)** *Fth1* and **(G)** *Ftl1* in primary neurons treated for 24 h with conditioned media (CM) from WT- or FpnKO-Microglia. **(H)** Expression of *Fth1* in primary neurons treated with CM from WT- or Fth1KO-Microglia. **(I)** Primary neuron expression of *Fth1* after 24 h exposure to CM from LD⁻ vs LD⁺ microglia. Points denote biological replicates; bars show mean ± s.e.m. Two-group comparisons used two-tailed unpaired Student’s t-tests. Significance is indicated as: *P ≤ 0.05, **P ≤ 0.01, ***P ≤ 0.001.

### Neuronal lipid droplet accumulation is also driven by iron accumulation

Lipid-droplet accumulation is a hallmark of neurons during aging and neurodegenerative diseases^2–5^. To model this in vitro, primary neuron were treated with oleic acid (OA; 50 µM) to drive LD synthesis. After 24 h, BODIPY staining revealed robust lipid accumulation (**Figure 6A**). A lower dose (25 µM) also increased lipid content, and after 48 h both OA concentrations produced a further rise in LD burden (**Figures S6A&S6B**). This lipid accumulation was accompanied by increased expression of Acc and Gpat1, supporting fatty-acid and triacylglycerol synthesis, respectively (**Figures 6B&6C**). Notably, LD accumulation in neurons failed to induce *Fth1* but upregulated *Fpn*, pointing to a shift toward iron export, possibly to prevent iron buildup and associated oxidative stress during lipid remodeling (**Figures 6D&6E**). When neuronal iron was further raised by adding FAC alongside OA, *Fth1* showed a slight increase, while *Fpn* surged by ∼10-fold, supporting a model in which neurons preferentially upregulate iron efflux rather than expand ferritin storage under lipid-loading and iron-rich conditions (**Figure S6C**). To probe the response mechanism under iron excess, we treated primary neurons with FAC alone. FAC increased lipid accumulation and reduced expression of pAMPK, a key regulator known to prevent lipid accumulation in neurons (**Figures 6F&6G**). Bulk RNA-seq of FAC-treated neurons showed a coherent transcriptional shift, with unsupervised PCA cleanly separating FAC from control samples along PC1 and tight replicate clustering (**Figure S6D**). Gene-set z-score summaries indicated coordinated remodeling of iron, lipid, and redox programs. In the iron module, iron import decreased, whereas storage/buffering and export/oxidation increased, including induction of *Ftl1* and *Fpn* (**Figure 6H**). Mitochondrial Fe-S biogenesis and iron homeostasis genes were elevated, as were heme catabolism/redox and ferroptosis defense components, consistent with adaptation to iron excess. In lipid pathways, neurons showed increased expression of lipid-droplet structural proteins, fatty-acid uptake and transport genes, pentose-phosphate/NADPH generators, and trafficking/LD-dynamics genes, matching the observed rise in neutral-lipid signal (**Figure 6I**). Redox programs were broadly induced, including superoxide dismutases, catalase/peroxiredoxins, glutathione and thioredoxin systems, NADPH-producing enzymes, NRF2 targets, and the CoQ/FSP1 arm, indicating reinforcement of antioxidant capacity under iron-rich conditions (**Figure 6J**). Because our microglia data indicated that iron-driven ROS is necessary for LD accumulation, we asked whether ROS scavenging could rescue neuronal LDs. NAC reduced FAC-induced lipid accumulation but did not reverse OA-only LDs (**Figure S6E**). Concordantly, NAC lowered *Acc* expression under OA+FAC, but not with OA alone (**Figure S6F**). Together, these data show that iron loading, but not OA alone, drives ROS-dependent LD accumulation in neurons and elicits an adaptive program characterized by reduced iron import, enhanced iron export and buffering, activation of LD biogenesis, and reinforced antioxidant defenses.

**Figure 6.**
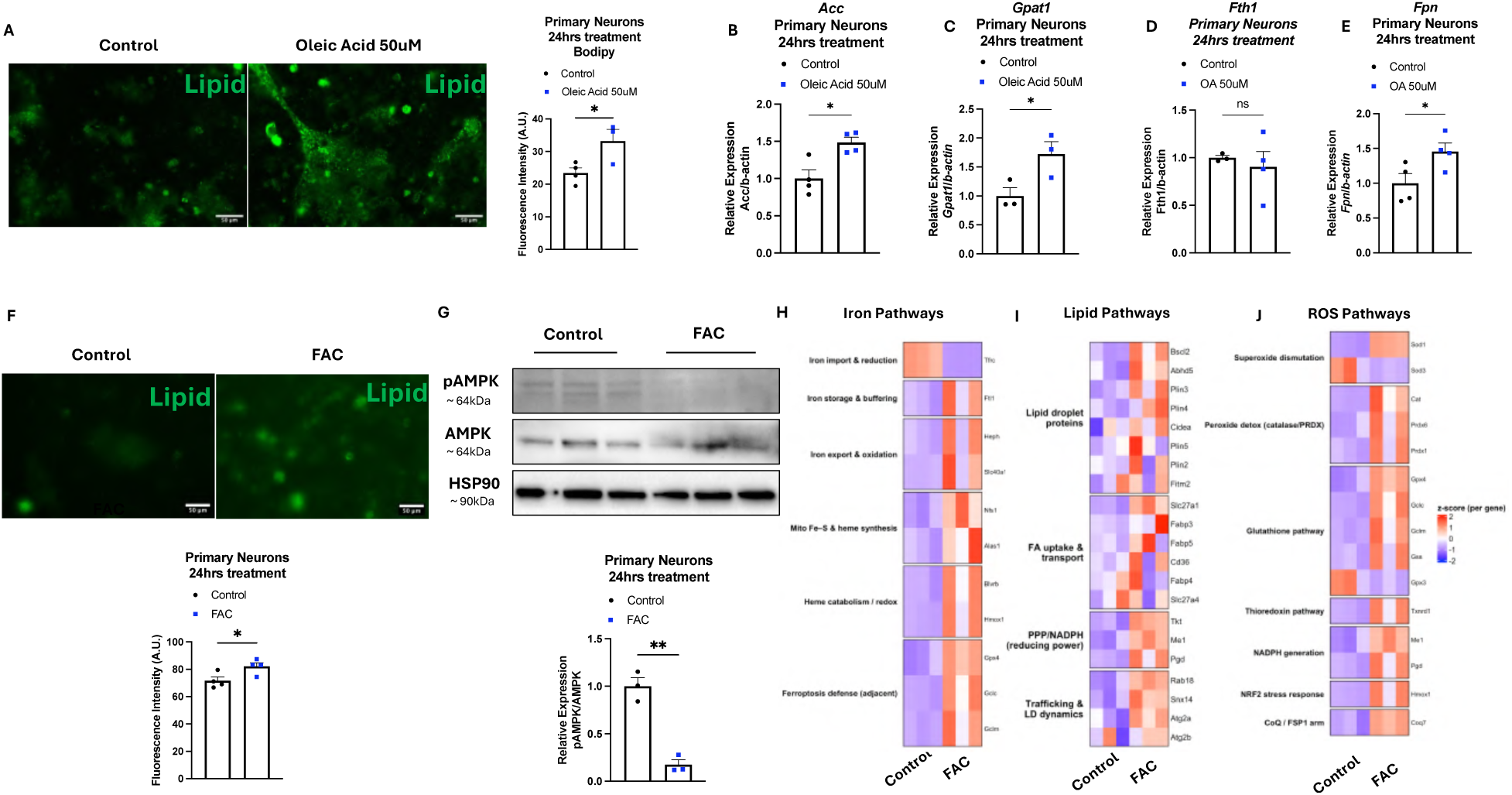
Iron accumulation drives ROS-dependent lipid-droplet accumulation and coordinated iron/lipid/redox transcriptional programs in neurons. **(A)** BODIPY staining images and fluorescence quantification (arbitrary units, A.U.) of primary neurons after 24 h treatment with oleic acid (OA, 50 µM) vs control. **(B-C)** Primary neurons expression of (B) *Acc*, and (C) *Gpat1* after 24 h of treatment with OA (50 µM) vs control. **(D-E)** Primary Neurons expression of (D) *Fth1* and (E) *Fpn* after 24 h treatment with OA (50uM). **(F)** BODIPY staining images and fluorescence quantification (arbitrary units, A.U.) of primary neurons after 24 h treatment with ferric ammonium citrate (FAC, 100 µM) vs control. **(G)** Immunoblot and its quantification of pAMPK, total AMPK, and HSP90 in control vs FAC-treated primary neurons. **(H–J)** Bulk RNA-seq pathway summaries (per-gene z-scores) for neurons treated 24 h with FAC vs control, showing **(H)** iron pathways, **(I)** lipid pathways, and **(J)** oxidative-stress programs. Points represent biological replicates; bars, mean ± s.e.m. Two-group comparisons used two-tailed unpaired Student’s t-test; when symbols are used: *P ≤ 0.05, **P ≤ 0.01. scale bars, 100 µm

## DISCUSSION

Here we show that aging drives a Fe2+-mitochondria axis in microglia that fuels ROS-dependent LD accumulation and reshapes microglia-neuron crosstalk. In the mouse hippocampus, aging increases the proportion of LD⁺ microglia with reduced mitochondrial mass, ferritin upregulation, and labile Fe²⁺ overload, accompanied by lipid peroxidation. LPS treatment increases Fe²⁺, mitochondrial ROS, and LDs in BV2 cells, whereas iron chelation or ROS scavenging blunts lipogenesis. Microglia-specific *Fth1* loss increases labile and mitochondrial Fe²⁺, raises ROS, and expands the LD⁺ fraction. Spatial and ligand-receptor analyses show reduced microglia-neuron distances and increased interactions with age, and conditioned media from LD⁺ microglia of aging mice or from microglia-specific Fth1-KO mice increases neuronal *Fth1* expression, indicating that iron-stressed microglia can reshape neuronal iron handling. Microglia-specific *Fpn* deletion minimally affects microglial Fe²⁺ yet lowers hippocampal FTH1, supporting a role for microglia in setting parenchymal iron availability. In neurons, iron excess preferentially induces iron export over storage and activates antioxidant and ferroptosis- mitigating programs while still increasing LD formation. NAC selectively reduces iron-driven LD formation. Together, these findings support a model in which aging reprograms microglia toward iron and ROS accumulation, promoting LD biogenesis and intensifying microglia-neuron interactions that alter iron availability and drive neuronal reprogramming toward iron, lipid, and redox responses.

Microglia shape brain aging and neurodegeneration, with age-dependent subsets that perform distinct functions^1,16,42,51–56^. LDAM arise with age and display impaired phagocytosis, elevated ROS, and increased pro-inflammatory cytokine release^1^. While inflammation, increased fatty-acid production, and oxidative stress have been implicated in LDAM formation^1^, our findings elevate iron overload and mitochondrial dysfunction as upstream determinants, defining a Fe^2+^- mitochondria-ROS pathway that drives microglial lipid accumulation. LDAM are transcriptionally distinct from other age-associated microglial populations, including disease-associated microglia (DAM) and neurodegenerative microglia (MGnD)^1,17^. They show partial overlap with lipid-associated macrophages (LAM), a pro-inflammatory macrophage state present in obese adipose tissue^1,56^. In obesity, iron overload promotes a shift in adipose-tissue macrophages toward LAM^57–59^. Similarly, here we show that, in the aging brain, increased iron favors a shift of microglia toward the LDAM state.

Iron is essential for brain function. To reach the brain, iron must cross the blood-brain barrier, although key transport mechanisms remain incompletely understood^60–62^. Most neural cell types express iron-handling machinery, including importers, ferritin for storage, and ferroportin for export, allowing intracellular iron to be tightly regulated by compensatory responses that stabilize Fe²⁺ levels^63^. Excess brain iron has been reported in mice and humans and is associated with aging and neurodegenerative disease^32,64–68^. This accumulation localizes to vulnerable cell types and regions, including neurons and microglia in the cortex and hippocampus, where neuropathological changes are prominent^60^. Although the causes of iron elevation are not fully defined, age-related blood-brain-barrier changes, neuroinflammation, and hepcidin-ferroportin regulation have been proposed to reduce ferroportin abundance and promote parenchymal iron accumulation^32,69–71^. Excess intracellular Fe²⁺ drives oxidative stress through the Fenton reaction, generating reactive oxygen species that damage lipids, proteins, and nucleic acids and can culminate in cell death^20,72^. With aging, microglial iron accumulation is accompanied by increased ROS and lipid peroxidation, a hallmark of ferroptotic stress. Prior work indicates that microglia are highly susceptible to ferroptosis relative to neurons and astrocytes and can trigger ferroptosis in neighboring cells, implicating microglia as key contributors to neurodegeneration^73,74^.

To dissect how iron handling shapes microglial phenotypes in vivo, we used two complementary mouse models. Microglia-specific deletion of *Fth1* removed the principal intracellular iron buffer and increased labile and mitochondrial Fe²⁺, elevated mitochondrial ROS, and expanded the LD⁺ microglial fraction. In contrast, microglia-specific deletion of *Fpn* reduced iron export while produced minimal changes in microglial labile Fe²⁺, consistent with compensatory adjustments in uptake and storage. These findings indicate that loss of buffering exerts a greater impact on microglial iron pools than loss of export. Aging is linked to reduced FPN, consistent with our microglia-specific *Fpn* KO model^32,75^. At the same time, aging is associated with higher *Fth1* expression, which may appear paradoxical relative to our Fth1 KO model^32,76^. In aging, iron influx and retention increase while export via FPN declines; ferritin is therefore induced to sequester excess iron, but storage can become saturated and ferritinophagy can release Fe²⁺ back to the cytosol^32,77^. As a result, the labile Fe²⁺ pool can remain elevated even when *Fth1* is high.

By contrast, loss of *Fth1* removes the compensatory buffer and directly increases labile iron, leading to mitochondrial dysfunction and oxidative stress, as shown in other cell types^78,79^. We next examined consequences for neurons using conditioned media from iron-stressed microglia. LDAM have been implicated in neurodegeneration through impaired phagocytosis and secretion of pro-inflammatory factors, such as IL-8, IL-6, and IL-1β, which can promote neuronal iron uptake and ferroptosis^1,74^. In line with these reports, conditioned media from Fth1- KO microglia and from LD⁺ microglia increased neuronal *Fth1* expression, consistent with higher neuronal iron accumulation. We then suggested a connection between iron accumulation and neuronal lipid storage. Only recently LD have been shown as an important player for health and disease, with LD accumulation in the brain being linked with oxidative stress and neurodegenerative diseases^80–82^. For example, in tauopathy models, LD accumulation is most evident in microglia and astrocytes, with minimal accumulation in neurons, whereas in PD substantia nigra, LDs have been observed in both microglia and dopaminergic neurons^83,84^. Here, we show lipid accumulation in neurons accompanied by increased lipogenesis under iron excess.

Limitations and future directions include several areas. First, the source and routes of iron entry into the aging brain require deeper mechanistic study. We focused on *Fth1* and *Fpn*, but other regulators, including hepcidin, transferrin-receptor pathways, and ferritin trafficking, likely contribute. Second, our work centers on microglia-neuron interactions. Astrocytes and oligodendrocytes also influence neuronal iron metabolism, including possible transfer of ferritin from to oligodendrocytes to neurons via extracellular vesicles or other related mechanisms^85,86^. Third, the active components in microglial conditioned media remain to be defined. Extracellular vesicles containing ferritin, iron, RNAs, or proteins, as well as soluble cytokines, are plausible mediators. Finally, we did not test neuron-to-microglia feedback. Iron overload and reduced pAMPK in neurons could secondarily promote microglial lipid accumulation. In tauopathy, preservation of neuronal pAMPK limits lipogenesis and lipophagy and may reduce lipid transfer to microglia^19^.

In summary, aging shifts microglia toward iron retention and mitochondrial stress that drive ROS-dependent LD biogenesis and intensify microglia-neuron crosstalk. Restoring ferritin-mediated buffering, normalizing iron flux, and increasing anti-ferroptotic capacity emerge as testable strategies to reduce LD burden and protect microglial and neuronal homeostasis in the aging brain.

## METHODS

### Animals

C57BL/6 (B6) mice were housed in a specific pathogen-free facility on a 12-h light/dark cycle at 22 °C with ad libitum access to water and standard chow (LabDiet 5001; 56.7% kcal carbohydrate, 29.8% kcal protein, 13.4% kcal fat). Both female and male mice were used at 6, 8, 36, 44, 75, and 94 weeks of age. B6 wild-type (WT) mice and the following strains were obtained from The Jackson Laboratory: Fpn^flox^ (129S-Slc40a1^tm2Nca^/J; #017790^87^), Fth1^flox^ (B6.129-Fth1^tm1.1Lck^/J; #018063), Tmem119-2A-CreERT2 (C57BL/6-Tmem119^em1(cre/ERT2)Gfng^/J; #031820), and Tau P301S (PS19) (B6;C3-Tg(Prnp-MAPT*P301S)PS19Vle/J; #008169).

Animals were acclimated for 1 week in the University of California, San Diego animal facility prior to experiments and breeding. To generate microglia-specific knockout mice, Fth1^flox/flox^ or Fpn^flox/flox^ mice were crossed to Tmem119-2A-CreERT2 mice, yielding Fth1- and Fpn- specific microglia KO, respectively. In Tmem119-2A-CreERT2 mice, Cre activity was induced by intraperitoneal tamoxifen at 75 mg/kg once daily for 5 consecutive days. All animal procedures were performed under UC San Diego guidelines for laboratory animal care and use, with random assignment of animals to cohorts.

### LPS in vivo treatment

Following what previous performed by Marschallinger et al., 2020^1^, with minor modifications, lipopolysaccharide (LPS; E. coli O111:B4; Sigma-Aldrich, Cat# L2630) was administered intraperitoneally at 1 mg/kg once daily for 3 days to 8-week-old wild-type (WT) mice. Sterile saline served as the vehicle control. Mice were euthanized 24 h after the final injection, and brains were collected for flow-cytometric analysis.

### BV2 cell culture and treatments

Murine BV2 microglia were maintained in high-glucose DMEM (4.5 g/L) supplemented with 10% FBS, 1% L-glutamine, and 1% penicillin-streptomycin at 37 °C in 5% CO₂. For experiments, cells were plated in 12- or 24-well plates and, at confluence, treated with LPS (5 µg/well) alone or in combination with ferric ammonium citrate (FAC; 100 µM; Sigma-Aldrich, Cat# F5879), N-acetylcysteine (NAC; 1 mM; Thermo Fisher Scientific, Cat# A15409.14), or 2,2′-bipyridine (Bipy; 50 µM; TCI America, Cat# B0468) for 24 h. After treatment, cells were stained with BODIPY 493/503 for lipid-droplet quantification and analyzed by flow cytometry or lysed for RNA isolation and qPCR analysis.

### Microglia Isolation

Microglia isolation followed published protocols with slight modifications^88,89^. Briefly, after intracardiac perfusion with ice-cold PBS, whole brains were rapidly removed into ice-cold PBS and mechanically meshed through 70-μm cell strainers to obtain a cell suspension. Cells were pelleted (600 x g, 10 min, 4 °C), resuspended in 6 mL 37% isotonic Percoll (Cytiva, Cat# 17089101), and carefully underlaid with 5 mL 70% isotonic Percoll in a 15-mL conical tube using a glass pipette. Gradients were centrifuged at 600 x g for 40 min at 16-18 °C with no acceleration and no brake. Mononuclear cells at the 37%/70% interphase were collected, diluted with ≥3 volumes of ice-cold PBS, and washed once (600 x g, 10 min, 4 °C). The final pellet was resuspended in staining buffer for flow cytometry analysis.

### Flow cytometry analysis and sorting

After microglia isolation, the cell pellet was resuspended in PBS with 2% FBS (optionally 2 mM EDTA) and incubated with Fc receptor block (anti-CD16/32; BioLegend, Cat# 101330; 1:200; 30 min; 4 °C). Surface staining was then performed in the dark for 30 min at 4 °C (or at RT when functional probes were included) with fluorochrome-conjugated antibodies to CD11b-BV421 (BioLegend, Cat# 101208; 1:200) and CD45-BV605 (BioLegend, Cat# 103122; 1:200) or CD45- APC-Cy7 (BioLegend, Cat# 103116; 1:200), together with LIVE/DEAD Fixable Aqua (Thermo Fisher, Cat# L34957; 1x). For selected experiments, in addition to the microglia panel, neutral lipids were labeled with LipidSpot 488 (Biotium, Cat# 70065; 1:1000, RT), and functional probes were applied immediately prior to acquisition: BioTracker Far-Red Labile Fe²⁺ Dye (Sigma-Aldrich, Cat# SCT037; 1:200; RT) for cytosolic labile Fe²⁺, mito-FerroGreen (Dojindo, Cat# M489; 1 µM; RT) for mitochondrial Fe²⁺, MitoTracker Green FM (Thermo Fisher, Cat# M7514; 100 nM; RT) for mitochondrial mass, and MitoSOX Red (Thermo Fisher, Cat# M36008; 5 µM; RT) for mitochondrial ROS. After antibody/dye staining, cells were washed once in PBS, filtered through a 40 µm strainer, and acquired/sorted on a Sony MA900 with single-stain and compensation controls. Microglia were gated as singlets → live → CD11b⁺ CD45^low^. Lipid- positive and -negative subsets were defined as CD11b⁺ CD45^low^ → LipidSpot⁺ and CD11b⁺ CD45^low^ → LipidSpot⁻, respectively. Full details of the gating strategy are provided in **Supplementary Materials (Supplementary Methods 1)**. Data were analyzed using FlowJo v10 software (BD Life Sciences).

### Primary neurons

Primary cortical/hippocampal neurons were isolated from C57BL/6J pups (Postnatal day 1-2) using a modified dissociation protocol^90^. Dissection solution (DS) was prepared by first making Solution A (137 mM NaCl, 5.4 mM KCl, 0.17 mM Na₂HPO₄, 0.22 mM KH₂PO₄ in ultrapure water) and Solution B (9.9 mM HEPES in ultrapure water), each stored at 4°C, then mixing 25 mL Solution A and 14 mL Solution B with 3 g D-glucose and 7.5 g sucrose, adjusting to pH 7.4, sterile-filtering (0.22 µm), and pre-chilling on ice. A trypsin inhibitor/BSA wash stock was prepared by dissolving 1 g trypsin inhibitor (Worthington, Cat# LS003087) and 1 g BSA in 20 mL DS (50 mg/mL each; pH 7.4), sterile-filtered, and kept on ice. Pups were rapidly decapitated, brains placed in ice-cold DS, cortices and hippocampi microdissected, meninges removed, and tissue minced on ice in DS. Minced tissue was transferred to a sterile dish and enzymatically dissociated in 5 mL pre-warmed 10X TrypLE Select (Thermo Fisher Scientific, Cat# A12177) for 25-30 min at 37 °C in the incubator, then collected into 15 mL conical tubes containing 10 mL ice-cold DS. Enzyme was quenched by five sequential washes: two “High Washes” prepared by mixing 600 µL trypsin inhibitor/BSA stock with 2.4 mL DS (3.0 mL total, divided into two 1.5 mL aliquots, used sequentially) followed by three “Low Washes” prepared by mixing 160 µL stock with 7.84 mL DS (8.0 mL total, divided into three ∼2.67 mL aliquots). For mechanical dissociation, three 1 mL pipette tips were cut to create decreasing bore sizes. Tissue was triturated 10-20 strokes with the widest tip, then with the intermediate tip, and finally with an uncut tip, keeping total trituration under 5 min and avoiding foaming. The cell-containing supernatant was transferred to a clean 15 mL conical, the dish was rinsed with 5 mL pre-warmed complete neuronal medium, and the rinse was combined for a final volume of 10 mL. Large tissue fragments were allowed to settle for ∼2 min, after which 9.5 mL of the cell suspension was carefully transferred to a new tube without disturbing debris. Twenty-four-well plates were coated with poly-D-lysine (50 µg/mL in PBS) for 1-2 h at room temperature, rinsed three times with sterile distilled water, and air-dried in a biosafety cabinet. Cells were plated in Neurobasal medium (Thermo Fisher Scientific, Cat# 21103049) supplemented with 2% B-27 (Thermo Fisher Scientific, Cat# A3582801), 1% GlutaMAX Thermo Fisher Scientific, Cat# 35050061), and 1% Antibiotic-Antimycotic (Thermo Fisher Scientific, Cat# 15240062). Cultures were maintained at 37 °C, 5% CO₂, with half-medium changes at 5-7 days, and experiments were performed on days 12-15 in vitro.

### Immunostaining to characterize primary neurons

To validate primary neuron cultures, neurons after 15 days in culture were fixed and stained for MAP2. Cultures were rinsed twice with PBS, fixed in freshly prepared 4% paraformaldehyde in PBS for 15 min at room temperature (RT), rinsed three times in PBS, permeabilized with 0.3% Triton X-100 in PBS for 5 min at RT, and rinsed three times in PBS. Non-specific binding was blocked with 5% goat serum (Thermo Fisher, Cat# 16210-064) in PBS for 60 min at RT. Primary antibody diluted in 5% goat serum was applied overnight at 4 °C (rabbit anti-MAP2, Thermo Fisher, Cat# PA5-17646; 1:100), followed by three 5-min washes in PBS. Secondary antibody was incubated for 60 min at RT in 5% goat serum (Alexa Fluor 488 goat anti-rabbit IgG [H+L], Thermo Fisher, Cat# A-11008; 1:200), then cells were washed three times in PBS. Nuclei were counterstained during the final wash with DAPI (3 ng/mL, 10 min), briefly rinsed in PBS, and imaged on a JuLI Stage fluorescence microscope (Nanoentek) using a 4x objective. Images were processed using ImageJ software (NIH) with a standardized deconvolution step for improved visualization.

### Microglia-conditioned media and neuronal exposure

Primary microglia were isolated as described and either FACS-sorted into LD⁺ and LD⁻ populations from aged mice or obtained from young microglia-specific *Fth1* and *Fpn* knockout mice and their WT littermates. Equal numbers of microglia were plated in 96-well plates (200 µL/well) in high-glucose DMEM (4.5 g/L) supplemented with 10% FBS, 1% L-glutamine, and 1% penicillin-streptomycin and maintained at 37 °C, 5% CO₂. After 24 h, conditioned media (CM) were collected, clarified to remove debris (300 X g, 5 min, 4 °C), aliquoted, and stored at - 80 °C until use. Primary neurons after 12-15 days in culture were exposed for 48 h to a 1:1 mixture of CM and neuron maintenance medium (200 µL CM + 200 µL neuron medium per well, 24-well). Following exposure, neurons were lysed in-well and RNA was isolated for qPCR as described.

### Primary neuron treatments

To induce lipid-droplet formation, primary neurons 12-15 days in culture, were treated with oleic acid (OA; Thermo Fisher Scientific, Cat# 031997.06) at 25 or 50 µM for 24 and 48 h. OA working solutions were prepared from a 5 mM OA-BSA stock generated by conjugating OA in 3% (w/v) fatty-acid-free BSA at 37 °C for 1 h, followed by 0.22 µm filtration; aliquots were stored at -20 °C and diluted into pre-warmed neuron medium immediately before use. For iron loading, FAC (Sigma-Aldrich, Cat# F5879) was applied at 100 µM, either alone or together with OA for the same exposure window. For ROS scavenging, NAC was included at 5 mM and added at the start of treatment. Experiments were performed in 24-well plates; for co-treatments (OA+FAC, OA+NAC, OA+FAC+NAC), reagents were added simultaneously. Media were prepared fresh, equilibrated to 37 °C/5% CO₂, and exchanged uniformly across conditions at the beginning of each treatment.

### BODIPY staining

Neutral lipids in BV2 cells and neurons were labeled with BODIPY 493/503 (Thermo Fisher Scientific, Cat# D3922). A 1 mg/mL stock was prepared in DMSO and stored at -20 °C, protected from light. For staining, BODIPY was diluted 1:1000 into culture medium and applied for 30 min at room temperature in the dark. Cells were rinsed two times with PBS and imaged immediately on a JuLI Stage fluorescence microscope (Nanoentek) using a 4x objective.

Fluorescence quantification was performed in ImageJ software (NIH) using identical parameters across conditions.

### Quantitative RT-PCR analysis

Total RNA was extracted from cells using TRIzol reagent (Invitrogen, Cat# 15596026) following the manufacturers’ instructions. Complementary DNA (cDNA) was generated with the High-Capacity cDNA Reverse Transcription Kit (Thermo Fisher Scientific, Cat# 4368813).

Quantitative PCR (qPCR) for mRNA targets was run in 10-µL reactions using PerfeCTa SYBR Green FastMix (Quantabio, Cat# 95073-05K) on a QuantStudio™ 3 real-time instrument.

Relative expression was calculated by the 2^−ΔΔCt^ method, normalizing mRNA to *Actb*, and values are reported as group means.

### Western Blot

Lysates from primary neurons or hippocampus were homogenized in RIPA buffer supplemented with protease and phosphatase inhibitors. Equal protein (5-10 µg per lane) was resolved by SDS– PAGE and transferred to membranes, which were blocked in 5% BSA/TBST and incubated overnight (4°C) with primary antibodies to FTH1 (rabbit, ABclonal, Cat# A1144, 1:1000), phospho-AMPKα (Thr183, Thr172) (rabbit, Thermo Fisher Scientific, Cat# 44-1150G, 1:1000), total AMPKα (rabbit, Thermo Fisher Scientific, Cat# PA5-116552, 1:1000), GAPDH (mouse, ABclonal, Cat# AC002, 1:1000), and HSP90 (mouse, Santa Cruz, Cat# sc-13119, 1:1000), followed by HRP-conjugated secondaries for 1 h at room temperature. Signals were developed using SuperSignal West Pico chemiluminescent substrate (Thermo Fisher Scientific, Cat# 34580) and imaged on a ChemiDoc XRS system (Bio-Rad) under exposure conditions verified to be within the linear range. Bands were analyzed in Image Lab v6.1 (Bio-Rad) and quantified in ImageJ software (NIH).

### Primary Neurons Bulk RNAseq

#### Library Preparation

Total RNA was extracted from primary neurons treated with vehicle (saline) or FAC (100 µM) using TRIzol, followed by cleanup with the Direct-zol RNA MicroPrep kit (Zymo, Cat# R2062) including on-column DNase I digestion (15 min, RT). RNA quantity and integrity were assessed by Qubit and Bioanalyzer; samples with RIN ≥ 7 were advanced. Poly(A)+ RNA was enriched (NEB, E7490) and strand-specific libraries were prepared with the NEBNext Ultra II Directional RNA Library Prep kit (NEB, E7760). Briefly, poly(A)+ RNA was fragmented at 94 °C for 15 min, first- and second-strand cDNA were synthesized, and double-stranded cDNA was purified with 1.8× SPRIselect. End repair/A-tailing and adaptor ligation (NEBNext Adaptor for Illumina) were followed by 0.9x SPRIselect cleanup. Libraries were PCR-amplified for 13 cycles with NEBNext Multiplex Oligos (NEB, Cat# E7335), size-selected at 0.8 x SPRIselect, pooled, and sequenced on an Illumina NovaSeq X Plus (paired-end 150 bp).

#### RNA-seq preprocessing

Paired-end RNA-seq libraries from primary neurons (control and FAC-treated) were processed using a standardized workflow. Raw FASTQ files underwent quality control with FastQC (v0.11.9) and report aggregation with MultiQC (v1.15). Adapters/low-quality bases were trimmed using fastp v0.23.4 (auto adapter detection, Q20 cutoff, minimum length 30 nt). Reads were aligned to mm39 (GRCm39) with STAR v2.7 (coordinate-sorted BAM, primary alignments only). Gene-level quantification used featureCounts (Subread v2.0.6) in paired-end, reverse-stranded mode against the matched GTF. Sample-by-gene counts were merged into a single matrix for downstream DE analysis.

#### RNA-seq Analysis

Program heatmaps (primary neurons). We started from DESeq2 results (Control vs FAC) and kept significant genes (padj < 0.05, |log₂FC| > 0.5). Normalized count matrices were loaded and, when needed, Ensembl IDs (version-stripped) were mapped to mouse symbols with org.Mm.eg.db, while unmapped genes were dropped. We curated gene panels a priori for (i) lipid/lipogenesis, (ii) iron biology (import, storage, export, Fe–S/heme, heme catabolism/redox, regulation, ferroptosis defense), and (iii) ROS/antioxidant systems (sources, dismutation, peroxide detox, glutathione/thioredoxin, NADPH, NRF2, CoQ/FSP1). For each panel, we intersected the curated list with the significant DE genes present in the matrix, extracted the submatrix, and Z-scored per gene across samples. Heatmaps were drawn with ComplexHeatmap (no clustering; blue-white-red scale), with group splits/titles and gene names shown.

### Bulk RNA analysis

Public bulk RNA-seq datasets were reanalyzed in R/RStudio using standard count-based workflows. For aging microglia (GSE208386^42^; mouse hippocampus, 3- vs 16-month), author-provided counts were modeled with DESeq2 (Wald tests on the age coefficient with apeglm LFC shrinkage); P values were adjusted by Benjamini-Hochberg, and DEGs were defined as padj < 0.05 with |log₂FC| > 1. For lipid-droplet status (GSE139542^1^; BODIPY^hi^/LD-high vs BODIPY^lo^/LD-low microglia), samples were annotated as LD-Low and LD-High and analyzed with DESeq2; to reduce noise/contamination we filtered baseMean ≤ 10 and removed a predefined neutrophil-gene list (Ngp, Ltf, S100a8, S100a9, Chil3, Lcn2, Camp, Cd177, Fcnb, Anxa1, Plbd1). DEGs were called at padj < 0.05 and |log₂FC| > 1. Pathway signature scores were also computed on GSE139542 by averaging DESeq2 size-factor-normalized counts across curated gene lists per sample, then comparing LD-High vs LD-Low with unpaired two-tailed t-tests (ggpubr). Gene sets were assembled a priori from the literature and included: ferroptosis (*Slc7a11*, *Gpx4*, *Ncoa4*, *Hmox1*, *Fth1*/*Ftl1*, *Slc40a1*, *Tfrc*, *Atg5*/*Atg7*, *Aifm2*/*Lpcat3*) and oxidative stress (*Nfe2l2*, *Nqo1*, *Hmox1*, *Gclc/Gclm*, *Sod1/2*, *Cat*, *Prdx1/2/6*, *Txn1/Txnrd1*, *Gpx1- 4*). For focused displays, enriched terms were filtered to mitochondria/oxidative-stress/lipid-metabolism pathways using keyword matching (e.g., “mitochond*”, “oxidative”, “lipid/lipogen*”, “fatty acid”, “peroxisome”, “ROS”, “beta-oxid*”). Visualizations (ggplot2/ggrepel/enrichplot) included PCA on VST counts (DESeq2 datasets), volcano plots (log₂FC vs -log₁₀ adjusted P; reference lines at |log₂FC| = 1, padj/FDR = 0.05), GO bar plots, and pathway-score box/jitter plots. Where indicated, gene-level box/jitter plots show DESeq2- normalized counts or VST values, with significance annotations reflecting the corresponding DESeq2 padj. All scripts export DEG tables, GO outputs, figures, and sessionInfo, which will be deposited on GitHub upon publication.

### Single-cell RNA and single-nucleus RNA sequencing analysis

We jointly analyzed mouse and human hippocampal 10x datasets: dentate gyrus (DG) scRNA-seq with matched Visium spatial data from C57BL/6 mice at 3 and 16-21 months (GSE233363^49^), mouse hippocampal sc/snRNA-seq at 4 and 24 months (GSE161340^44^), and human hippocampal snRNA-seq (GSE185553^43^, GSE199243^43^). Raw matrices were imported into Seurat v5, gene symbols were de-duplicated, and cohort-specific QC was applied (DG scRNA-seq: nFeature_RNA >500 & <6,000, nCount_RNA <30,000, percent.mt <15%; Ogrodnik scRNA-seq: 500-6,000 genes, 1,000-25,000 UMIs, ≤12% mtRNA; snRNA-seq: nFeature_RNA >200 & <6,000, percent.mt ≤5%; integrated human objects further filtered to nFeature_RNA >200 & <5,000, percent.mt <10%). DG scRNA-seq and human datasets were normalized with SCTransform v2 and integrated using SCT anchors (3,000 features); Ogrodnik sc/snRNA-seq used log-normalization, 3,000 variable features, and RPCA integration; for snRNA-seq in Seurat v5, normalized layers were merged to a single data matrix before subsetting/DE. Objects were reduced by PCA, neighbors computed, embedded by UMAP, and clustered across resolutions. Broad identities (Neuron, Microglia, Other) were assigned by canonical markers (neurons: *Snap25*/*Rbfox3*/*Syt1*/*Map2*/*Tubb3*, and microglia: *P2ry12*/*Tmem119*/*Cx3cr1*/*Csf1r*/*C1qa*). Young vs Aging comparisons (mouse: 3 vs 16-21 months; 4 vs 24 months; human: 18-39 vs 60-95 years) were performed within neurons and microglia using Seurat’s Wilcoxon test (min.pct ≥0.1, |log2FC| >0.25 unless noted), excluding mitochondrial genes prior to Benjamini-Hochberg correction (padj <0.05). Within the microglia subset only, we computed per-cell log-normalized expression for *Fth1*. Cells were then split into quartiles by their *Fth1* value (Fth1-High, top quartile and Fth1-Low: bottom quartile). GO-BP enrichment used clusterProfiler (org.Mm.eg.db for mouse; org.Hs.eg.db for human), and iron/ferroptosis programs (e.g., *Fth1*/*Gpx4*/*Slc7a11*) were profiled at single-cell and pseudobulk levels. Neuron-microglia signaling was inferred with CellChat (CellChatDB.mouse) in Young vs Aging groups. For Visium, neuron-to-microglia distances were computed as nearest-neighbor spot-to-spot (FNN), converted to micrometers with slide-specific scale factors, and compared by Wilcoxon tests. All scripts export DEG tables, GO outputs, figures, and sessionInfo, which will be deposited on GitHub upon publication.

### Statistical analysis

Mice were randomly assigned to experimental groups for all in vivo studies. Unless noted, data are shown as mean ± SEM. Comparisons between two groups used unpaired, two-tailed Student’s t-tests; analyses with more than two groups used one-way ANOVA with an appropriate post hoc test as indicated in the figure legends. RNA-seq analyses followed established pipelines with Benjamini-Hochberg multiple-testing correction. All analyses were performed in GraphPad Prism 10 (GraphPad Software) and RStudio; P ≤ 0.05 was considered statistically significant, with exact P values and significance symbols reported in the figure legends.

### Data and code availability

Public datasets reanalyzed include mouse dentate gyrus scRNA-seq with matched Visium spatial hippocampus (GSE233363), mouse hippocampus sc/snRNA-seq (GSE161340), human hippocampus snRNA-seq (GSE185553, GSE199243), bulk RNA-seq of mouse hippocampal microglia (GSE208386), and bulk RNA-seq comparing LD-high vs LD-low microglia (GSE139542). Bulk RNA-seq from primary neurons (FAC vs control) will be deposited in GEO upon publication. Custom code for preprocessing, Seurat/CellChat analyses, spatial nearest-neighbor distance calculations, and bulk RNA-seq (DESeq2/clusterProfiler) will be released on GitHub at publication. Additional data supporting this study are available from the Lead Contact upon reasonable request.

## Acknowledgments

This study was funded by National Institutes of Health grants (R01DK125560 to W.Y., R01AG074273 and R01AG078185 to X. C., and U24DK132746-01 to K.C.R.) and by the Larry L. Hillblom Foundation (2023-D-011-FEL to K.C.R.). This publication includes data generated at the UC San Diego IGM Genomics Center utilizing an illumina NovaSeq X Plus that was purchased with funding from a National Institutes of Health SIG grant (#S10 OD026929).

## Author contributions

Conceptualization: WYing, KCR

Methodology: WYing, KCR, XC

Investigation: WYing, KCR, QX, CQ, LW, WYuan, VB, YD, LV, DA, GA, JP

Visualization: WYing, KCR

Funding acquisition: WYing, KCR, WL

Project administration: WYing

Supervision: WYing, WL

Writing - original draft: WYing, KCR

Writing - review & editing: WYing, KCR, XC

---

**Figure S1.**
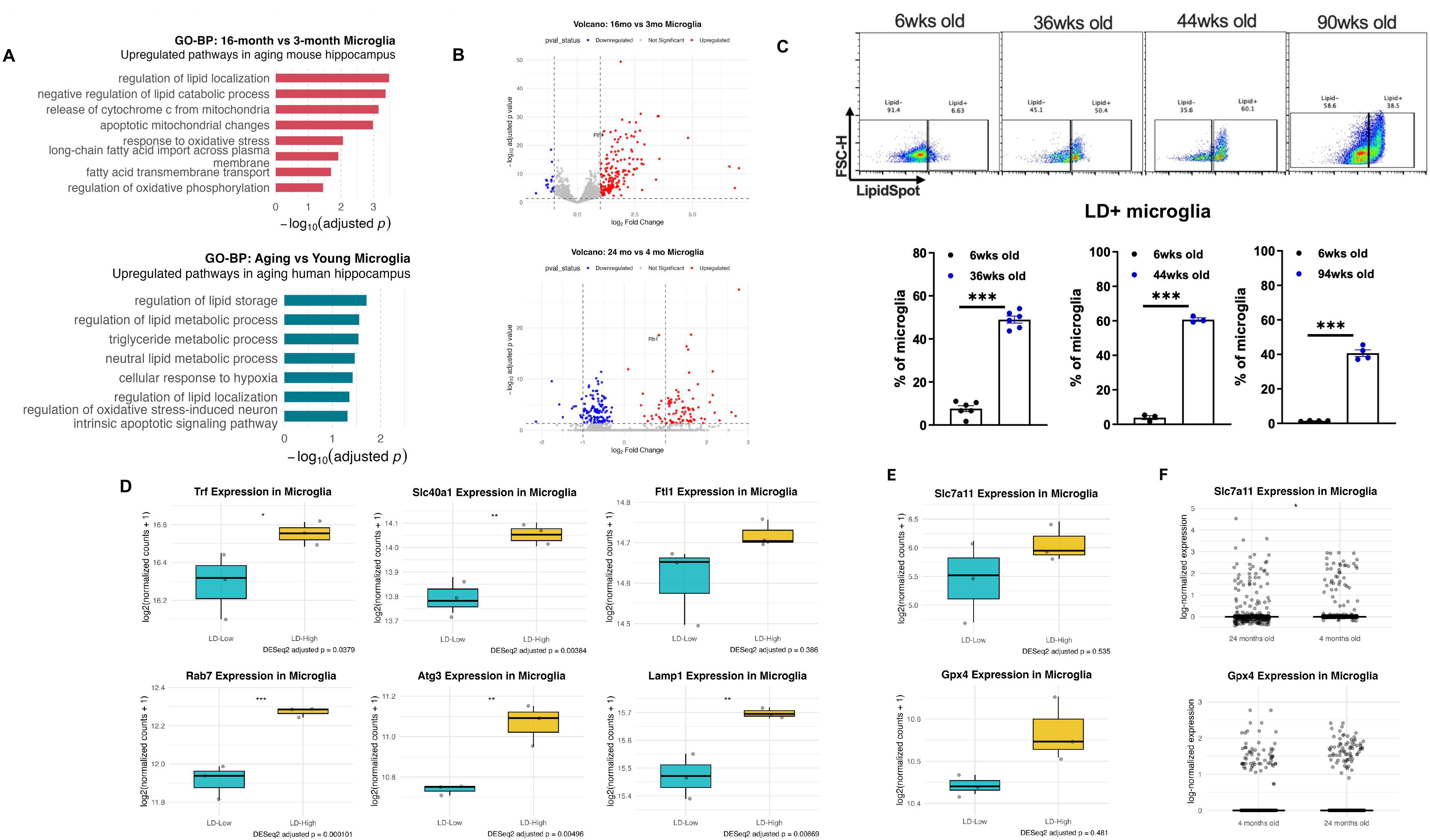
Microglial profiling across age and lipid-droplet status.

**Figure S2.**
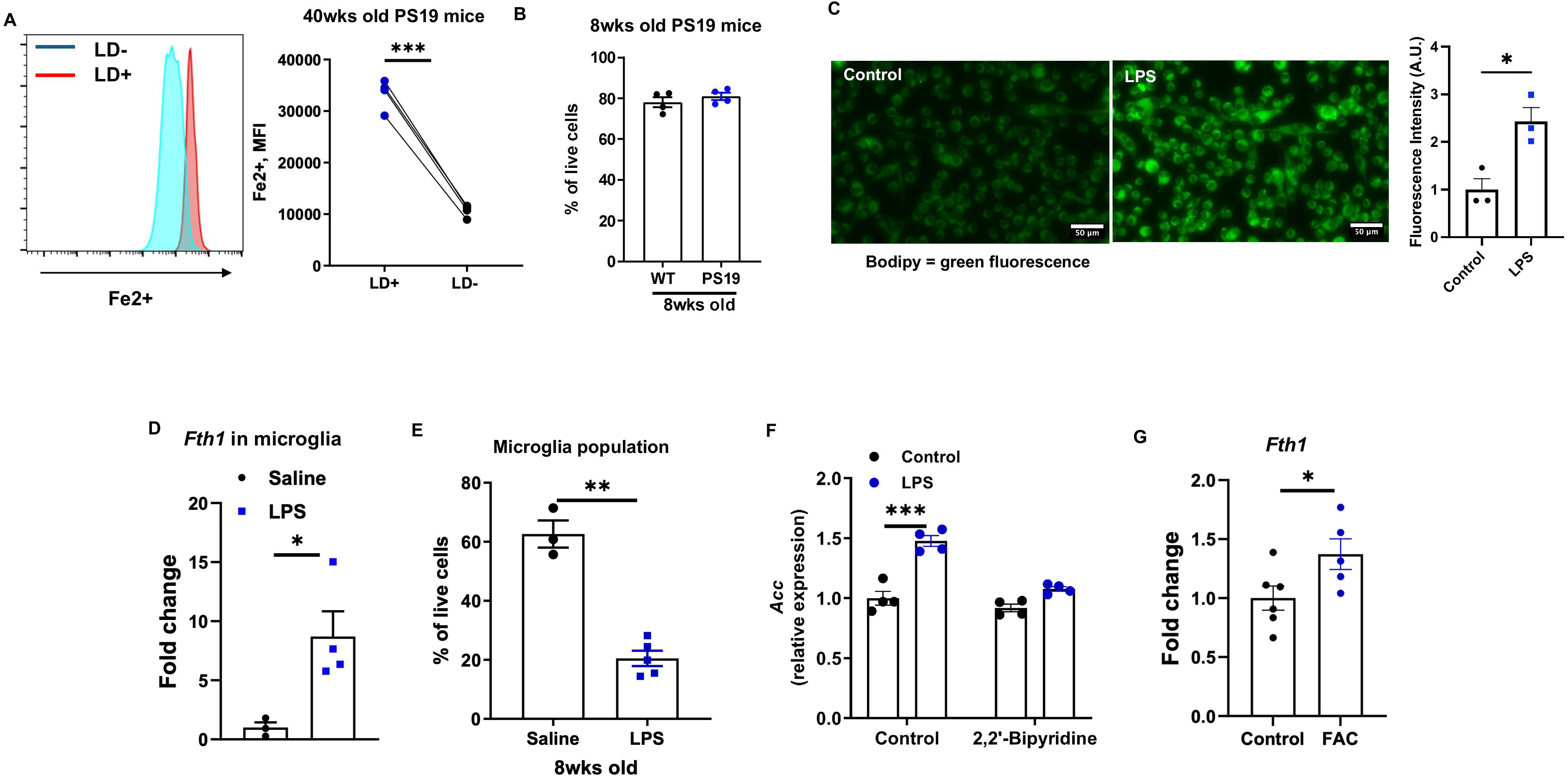
Iron status and lipid-droplet accumulation in aging microglia and LPS-treated BV2 cells.

**Figure S3.**
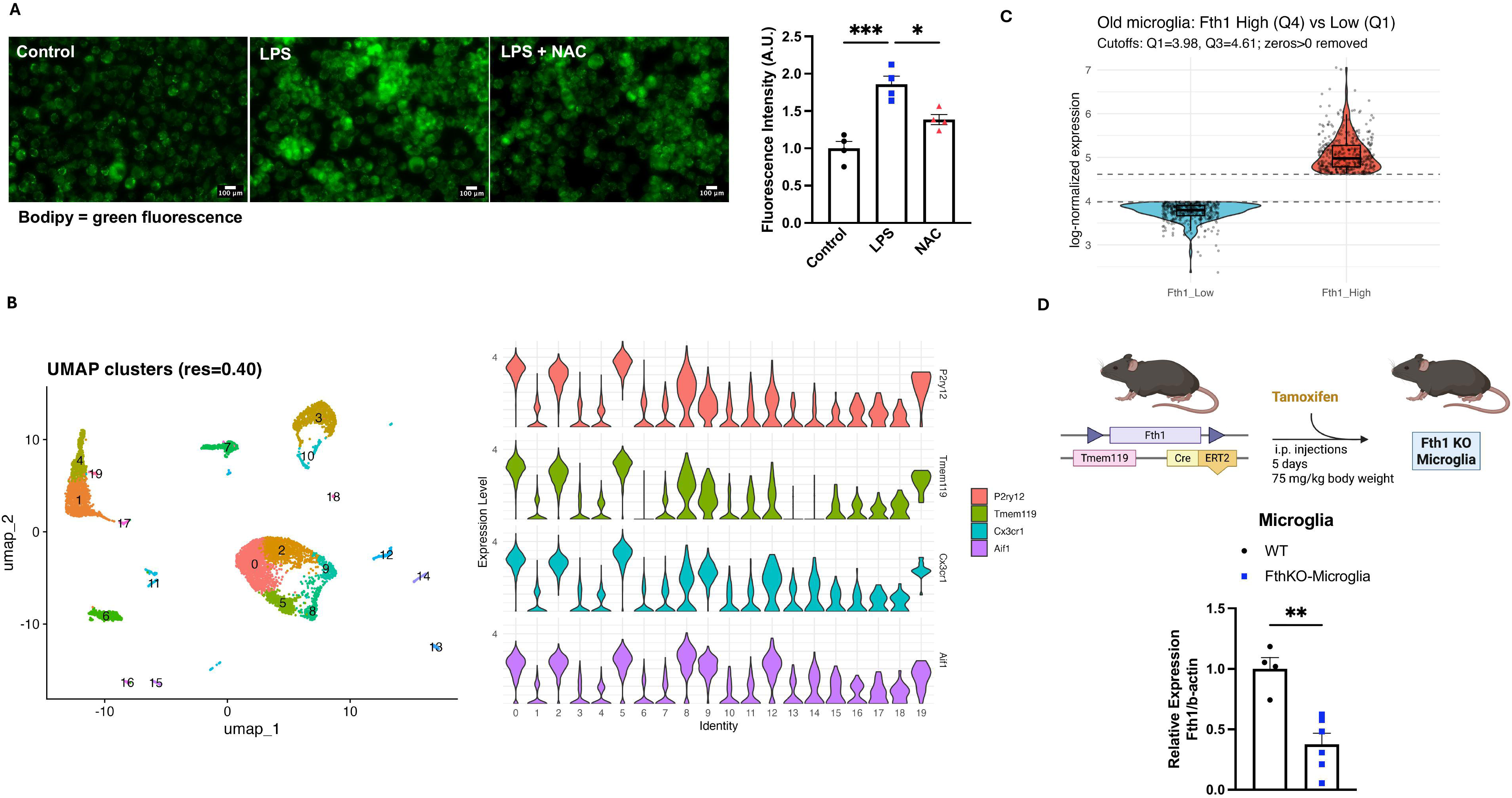
ROS modulation and LD assays in BV2, microglia scRNA-seq reference panels, and the Fth 1 KO model.

**Figure S4.**
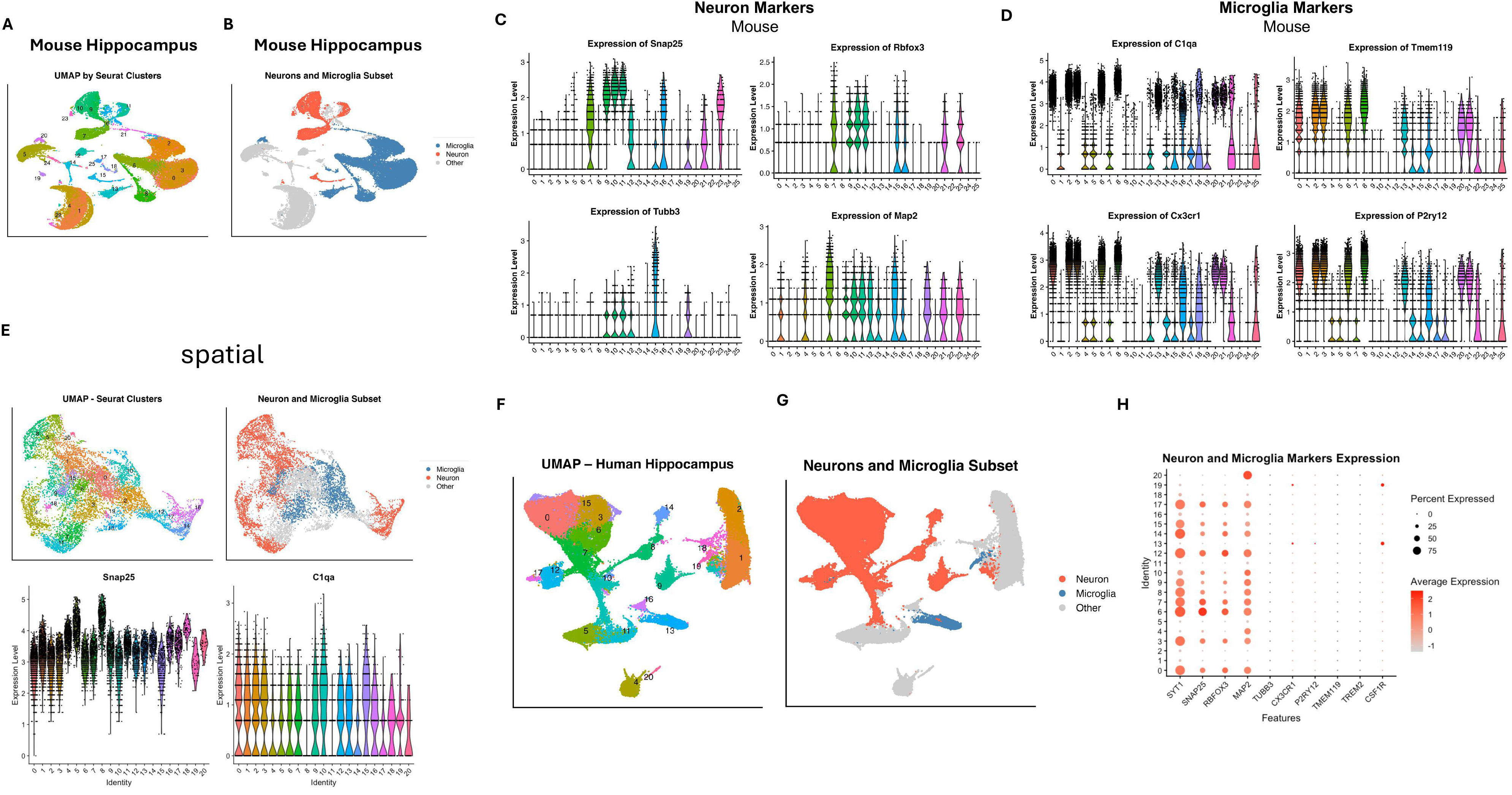
Cell-type validation across mouse and human hippocampus.

**Figure S5.**
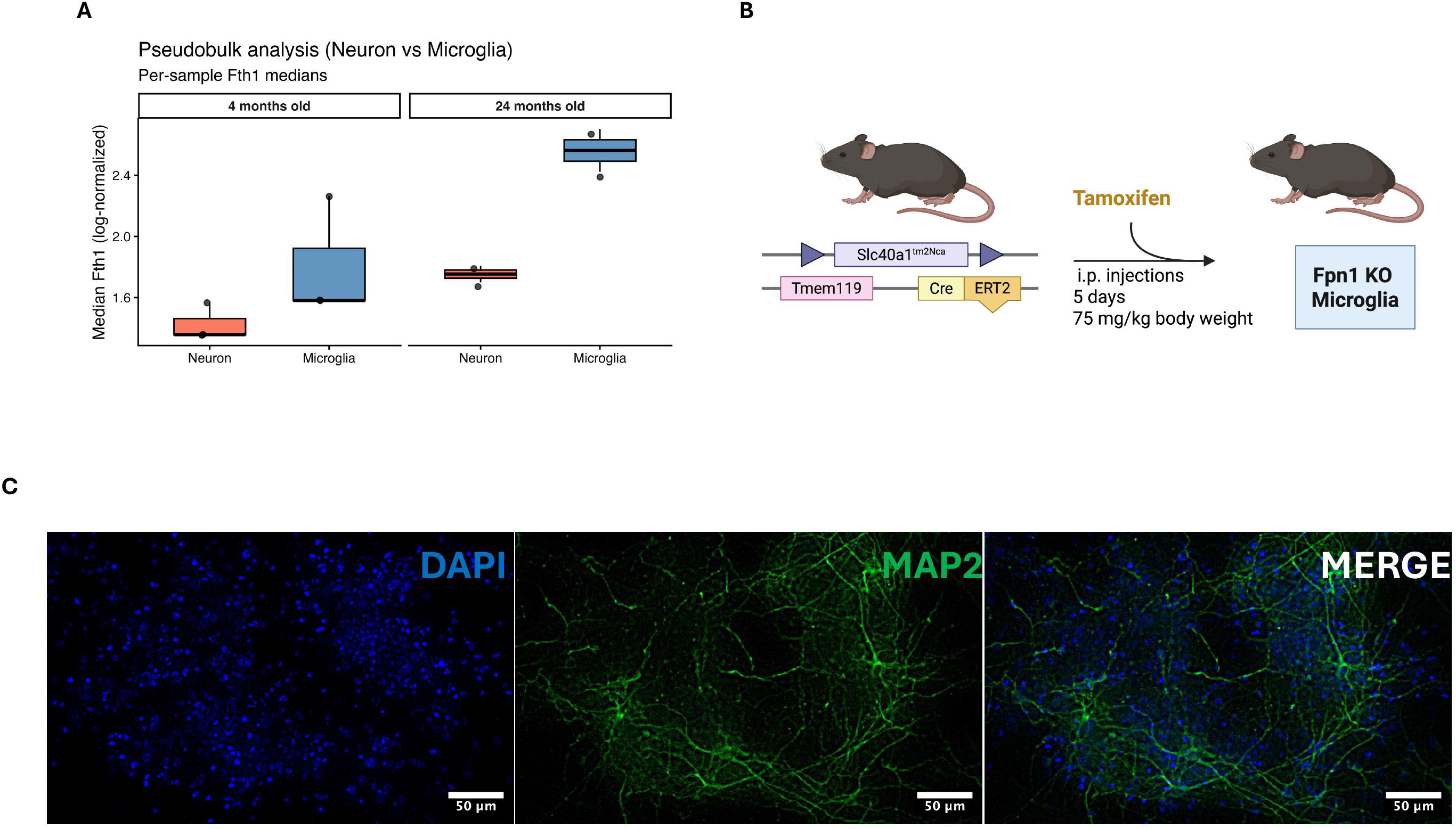
Pseudobulk ferritin, microglial FpnKO model, and primary-neuron validation.

**Figure S6.**
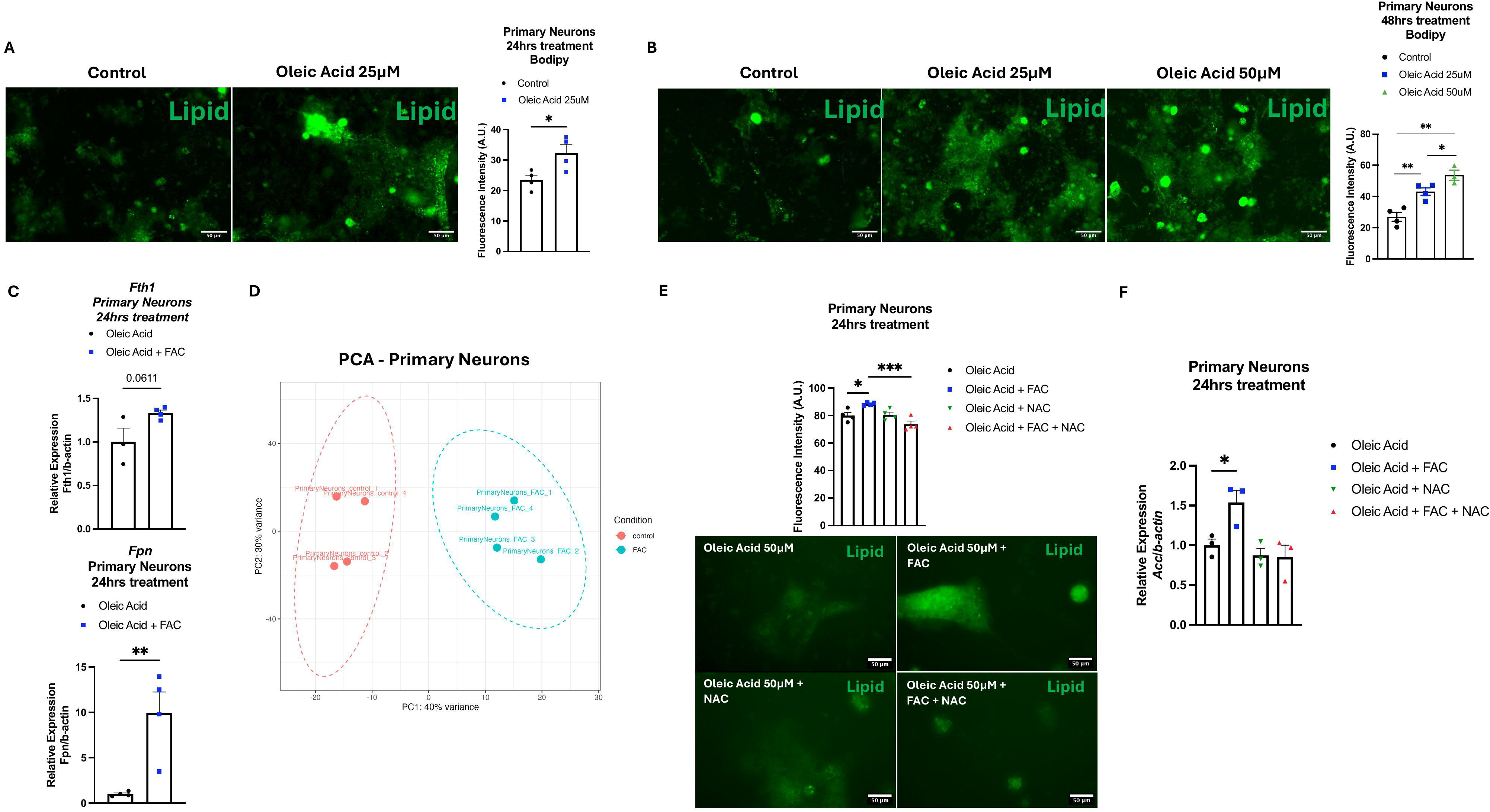
Dose-time responses and ROS control of neuronal LDs.

