## Supplementary Materials for "Labile iron overload reprograms microglia and neurons for lipid droplet synthesis in the aging brain"

### LIST OF SUPPLEMENTARY MATERIALS

#### Supplementary Methods:

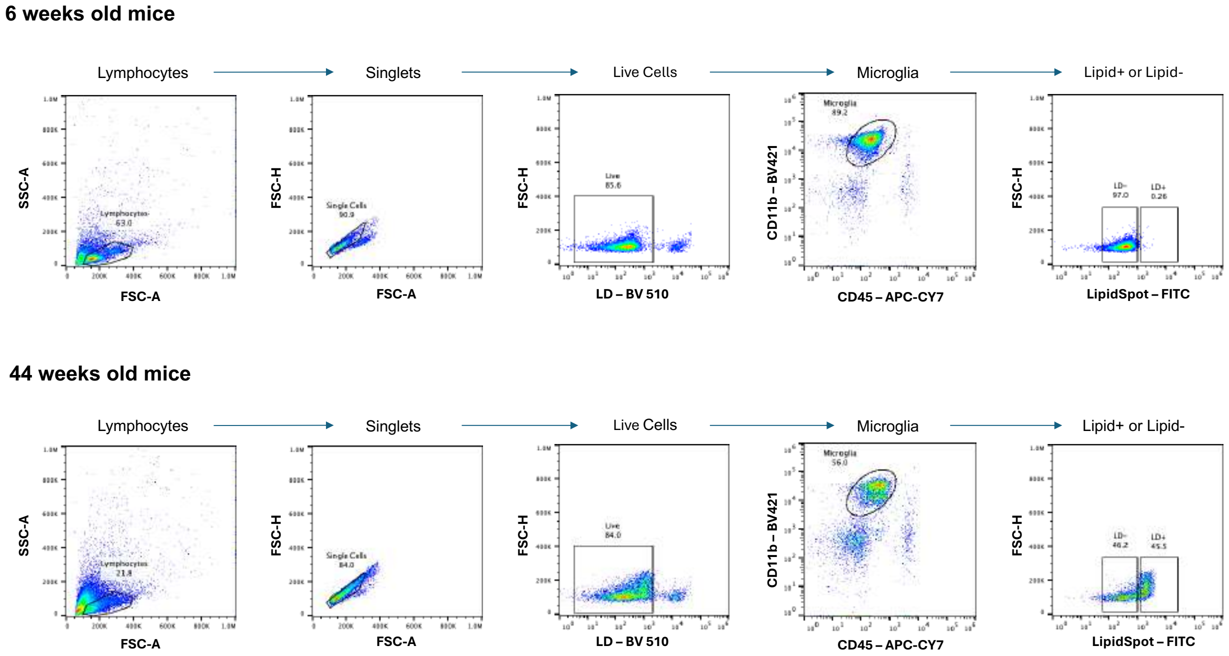


**Supplementary Methods 1. Gating Strategy for Microglia Lipid+ and Lipid- sorting.** Cells were first plotted on FSC-A vs SSC-A to exclude debris and place a broad leukocyte gate. Doublets were removed using FSC-H vs FSC-, followed by viability gating (LIVE/DEAD Aqua$^{-}$) to keep live singlets only. Microglia were identified as CD11b$^{+}$ CD45$^{low}$. Within this CD11b$^{+}$ CD45$^{low}$ gate (microglia), neutral-lipid status was selected with LipidSpot 488: Lipid$^{+}$ = LipidSpot$^{high}$ and Lipid$^{–}$ = LipidSpot$^{low/neg}$. Data were analyzed in FlowJo v10.

#### Supplementary Figures:

Figure S1. Microglial profiling across age and lipid-droplet status

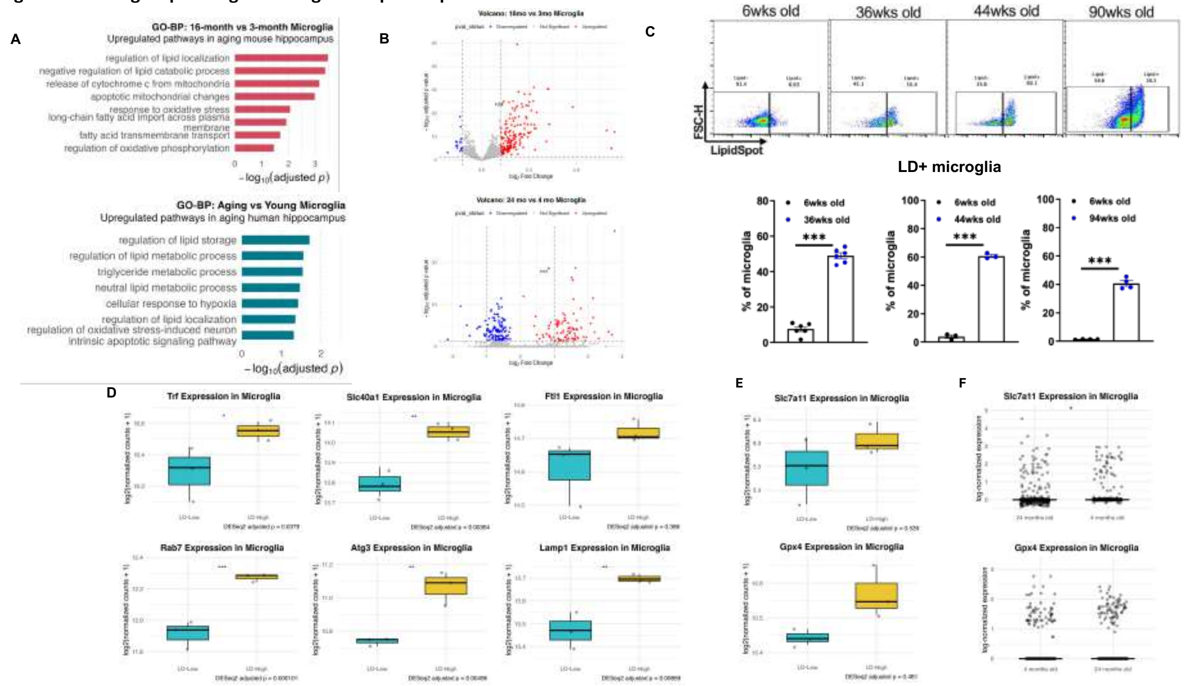


**Figure S1. Microglial profiling across age and lipid-droplet status. (A)** GO Biological Process (GO-BP) enrichment for upregulated genes in mouse hippocampal microglia (16 vs 3 months; top) and human hippocampal microglia (aging 60-95 years vs young 18-39 years; bottom). **(B)** Volcano plots of differential expression in mouse microglia (16 vs 3 months, top; 24 vs 4 months, bottom). **(C)** Top: flow-cytometric gating to define lipid-droplet-positive ($LD^+$) and $LD^-$ microglia using LipidSpot 488 within the $CD11b^+CD45^{low}$ gate. Bottom: percentage of $LD^+$ cells among whole-brain microglia at 6, 36, 44, and 94 weeks. **(D)** Bulk RNA-seq normalized counts for *Trf*, *Slc40a1*, *Ftl1*, *Rab7*, *Atg3*, and *Lamp1* in LD-high vs LD-low microglia isolated from 18-month mouse hippocampus. **(E)** Bulk RNA-seq normalized counts for *Slc7a11* and *Gpx4* in LD-low vs LD-high microglia. **(F)** scRNA-seq expression of *Slc7a11* and *Gpx4* in mouse hippocampal microglia (4 vs 24 months). Points represent biological replicates; bars, mean ± s.e.m. Unless indicated on panels: $*P \leq 0.05$, $**P \leq 0.01$, $***P \leq 0.001$. Two-tailed unpaired Student's t-test was performed on **(C)**. RNA-seq analyses (**D–E**) by DESeq2 with Benjamini-Hochberg FDR correction.

Figure S2. Iron status and lipid-droplet accumulation in aging microglia and LPS-treated BV2 cells

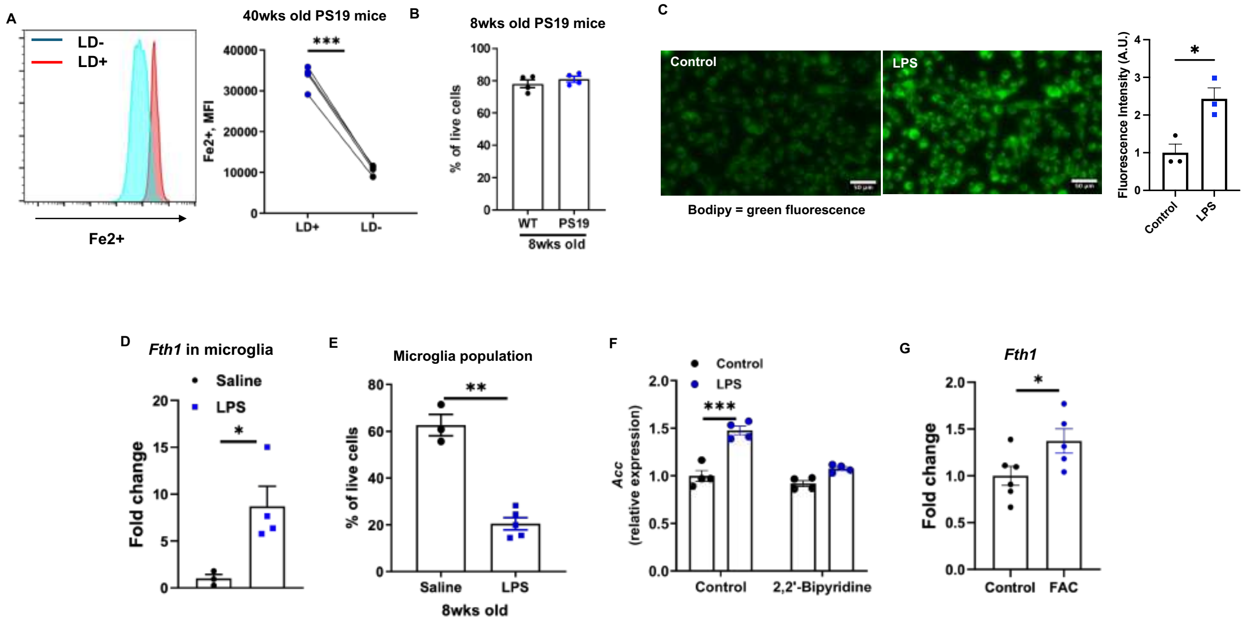


**Figure S2. Iron status and lipid-droplet accumulation in aging microglia and LPS-treated BV2 cells. (A)** Labile $Fe^{2+}$ mean fluorescence intensity (MFI) in $LD^{+}$ vs $LD^{-}$ microglia isolated from 40-week-old PS19 mice (LD status defined by LipidSpot). **(B)** Frequency of microglia ($CD11b^{+}CD45^{low}$) among live brain cells in 8-week-old WT vs PS19 mice. **(C)** BODIPY fluorescence images of BV2 cells after 24 h control (saline) or LPS (5 µg), with quantification (arbitrary units, A.U.). **(D)** Expression of *Fth1* in microglia sorted from saline- vs LPS-treated mice (1 mg/kg once daily for 3 days). **(E)** Frequency of microglia among live brain cells after saline vs LPS injections in 8-week-old mice (1 mg/kg once daily for 3 days). **(F)** Expression of *Acc* in BV2 cells after 24 h treatment with control vs LPS (5 µg) in the absence or presence 2,2′-bipyridine (Bipy, 50 µM). **(G)** Expression of *Fth1* in BV2 cells treated 24 h with ferric ammonium citrate (FAC, 100 µM) vs control. Points represent biological replicates; bars, mean ± s.e.m. Two-group comparisons: two-tailed unpaired Student's t-test. Multi-group comparisons: one-way ANOVA with Tukey's multiple-comparisons test. Exact P values are reported on plots; when symbols are used: $^{*}P \leq 0.05$, $^{**}P \leq 0.01$, $^{***}P \leq 0.001$. Scale bar, 100 µm.

Figure S3. ROS modulation and LD assays in BV2, microglia scRNA-seq reference panels, and the Fth1 KO model

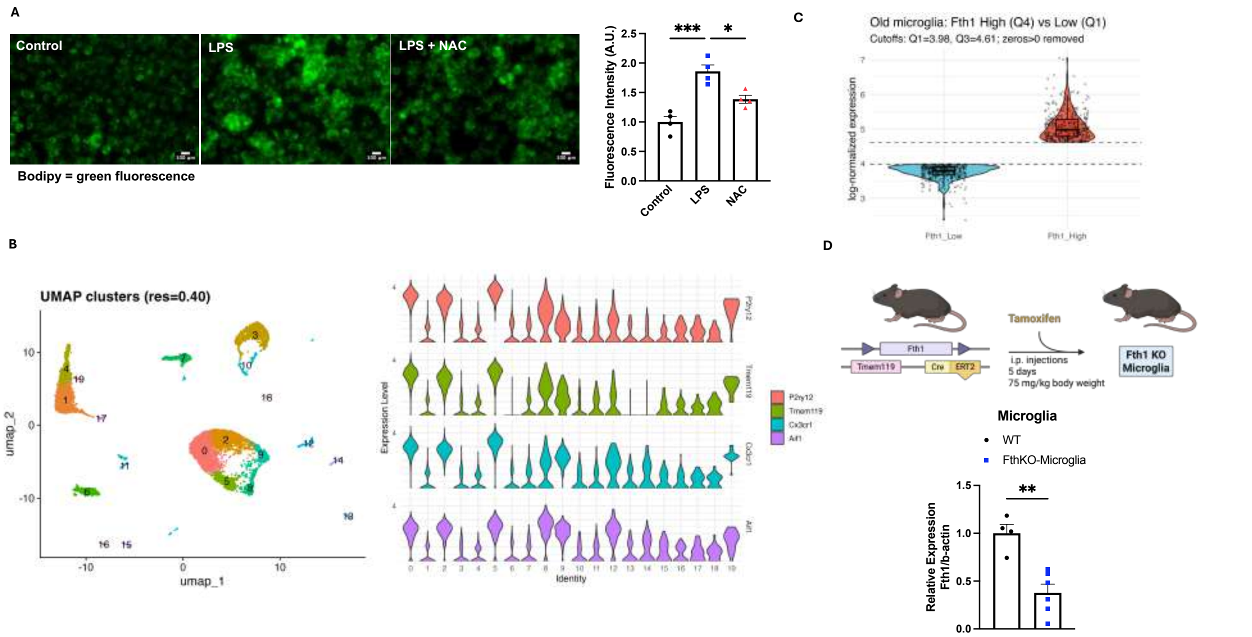


**Figure S3. ROS modulation and LD assays in BV2, microglia scRNA-seq reference panels, and the Fth1 KO model. (A)** BODIPY images and fluorescence quantification (arbitrary units, A.U.) of BV2 cells after 24 h treatment with Control (saline), LPS (5 µg), or LPS + N-acetylcysteine (NAC, 1mM). **(B)** scRNA-seq re-analysis of 24-month-old mouse hippocampus from: UMAP of clusters and violin plots for canonical microglial markers (*P2ry12*, *Tmem119*, *Cx3cr1*, *Aif1*) used to annotate the microglial population. **(C)** Differential expression summary within aging microglia stratified by ferritin heavy chain (*Fth1*) abundance (quartile 4, Fth1-High, vs quartile 1, Fth1-Low); distribution of log-normalized expression change shown per cell. **(D)** Schematic of the tamoxifen-inducible Tmem119-CreERT2/Fth1$^{fl/fl}$ microglia-specific knockout mouse model and qPCR validation of *Fth1* reduction in sorted Fth1KO microglia relative to WT littermates. Points represent biological replicates; bars, mean ± s.e.m. Two-group comparisons used two-tailed unpaired Student's t-test; multi-group comparisons used one-way ANOVA with Tukey's post hoc test. *P ≤ 0.05, **P ≤ 0.01, ***P ≤ 0.001. Scale bars, 100 µm.

Figure S4. Cell-type validation across mouse and human hippocampus

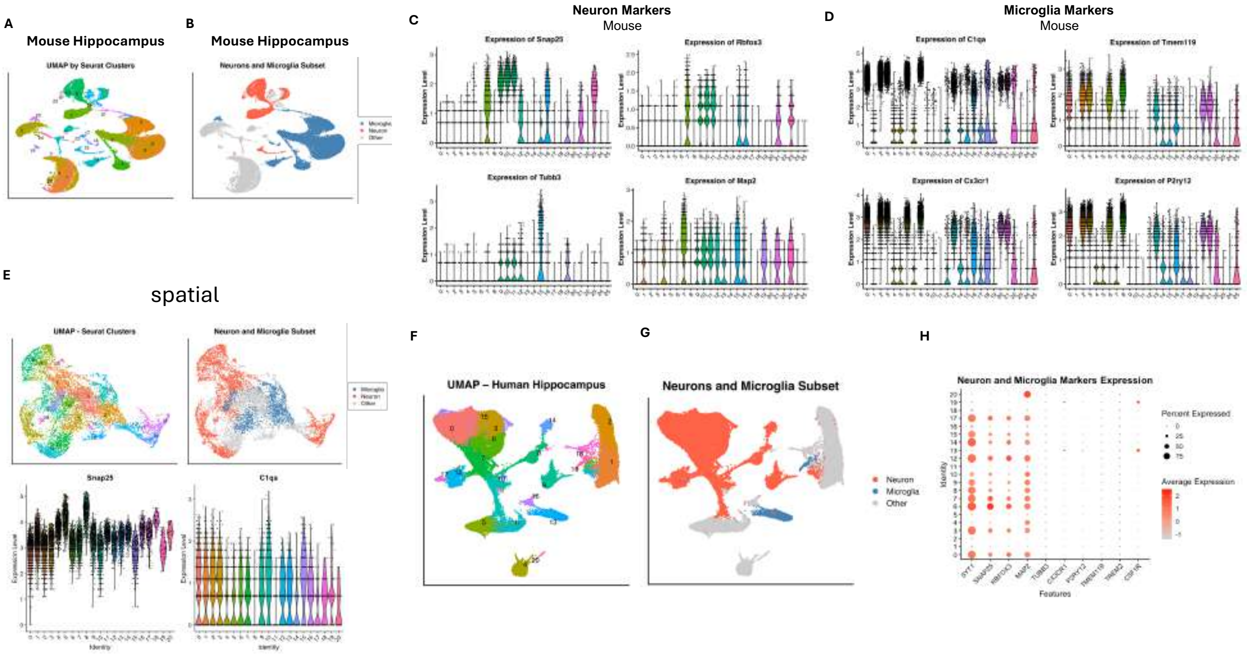


**Figure S4. Cell-type validation across mouse and human hippocampus. (A)** UMAP of mouse dentate gyrus scRNA-seq showing Seurat clusters. **(B)** UMAP view restricted to cell classes (Neuron, Microglia, Other) assigned by canonical markers. **(C)** Violin plots for neuronal markers (*Snap25*, *Rbfox3*, *Tubb3*, *Map2*) across mouse dentate gyrus scRNA-seq clusters. **(D)** Violin plots for microglial markers (*C1qa*, *Tmem119*, *Cx3cr1*, *P2ry12*) across mouse dentate gyrus scRNA-seq clusters. **(E)** Visium spatial mouse hippocampus showing spot-level UMAP/clusters, a Neuron/Microglia subset, and example marker expression (*Snap25*, *C1qa*). **(F)** UMAP of all clusters after human hippocampus snRNA-seq integration. **(G)** Human hippocampus snRNA-seq Neuron/Microglia subset on UMAP. **(H)** Dot plot confirming human neuronal (SYT1, SNAP25, RBFOX3, MAP2) and microglial (C1QA, TMEM119, P2RY12, CX3CR1) markers across integrated hippocampus snRNA-seq clusters.

**Figure S5. Pseudobulk ferritin, microglial FpnKO model, and primary-neuron validation**

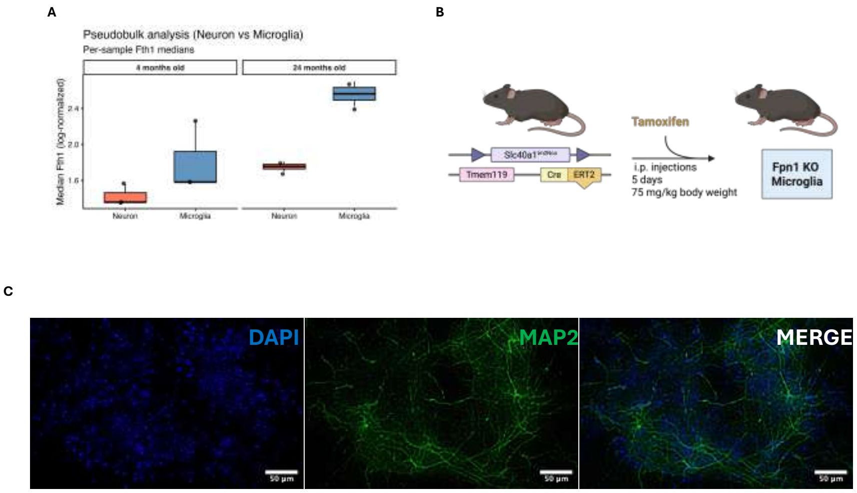


**Figure S5. Pseudobulk ferritin, microglial FpnKO model, and primary-neuron validation. (A)** snRNA-seq pseudobulk *Fth1* expression comparing neurons and microglia at 4 vs 24 months; boxes show median and IQR. **(B)** Schematic of the tamoxifen-inducible, microglia-specific ferroportin knockout (Tmem119-CreERT2/ Slc40a1$^{flox/flox}$). **(C)** Primary cortical/hippocampal neurons stained for MAP2 with DAPI nuclear counterstain; scale bars, 100 µm.

Figure S6. Dose-time responses and ROS control of neuronal LDs

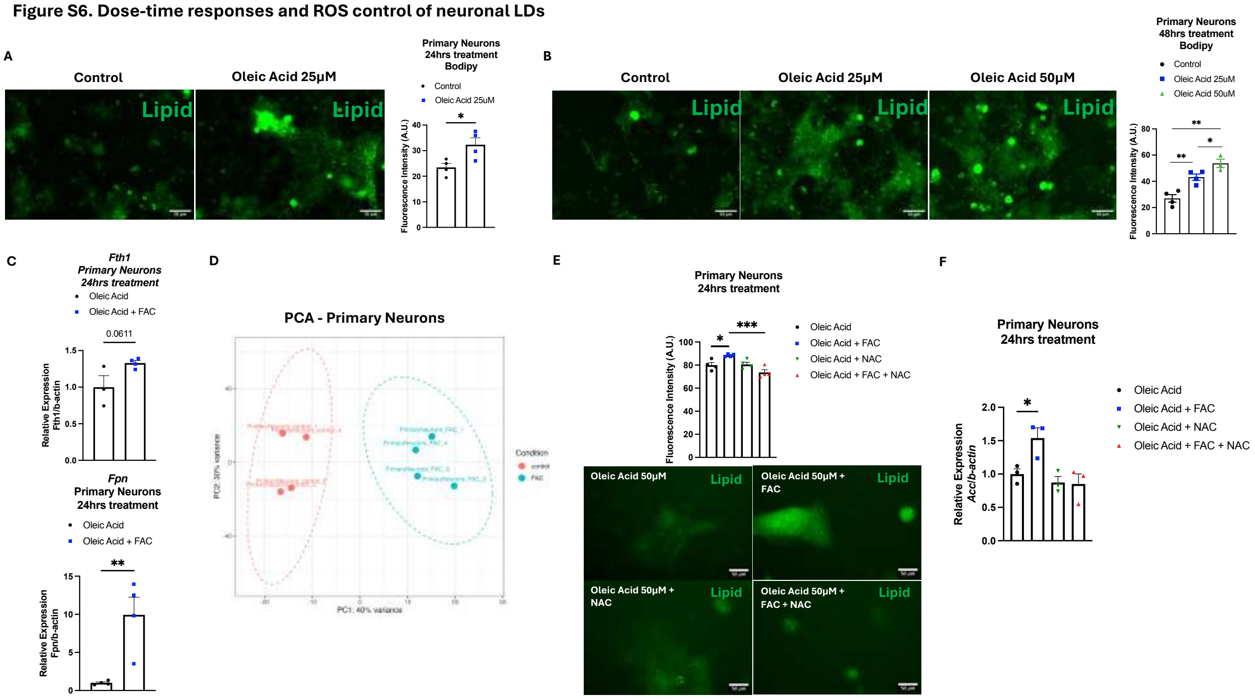


**Figure S6. Dose-time responses and ROS control of neuronal LDs. (A)** BODIPY staining images and fluorescence quantification (arbitrary units, A.U.) of primary neurons after 24 h oleic acid (OA, 25 μM) versus control. **(B)** BODIPY staining images and fluorescence quantification (arbitrary units, A.U.) of primary neurons treated for 48 h with control, OA 25 μM, or OA 50 μM. **(C)** Neuronal *Fth1* and *Fpn* expression after 24 h treatment of OA 50 μM with or without ferric ammonium citrate (FAC, 100 μM). **(D)** PCA of primary-neuron bulk RNA-seq showing separation of treatment groups. **(E)** BODIPY staining images and fluorescence quantification (arbitrary units, A.U.) of primary neurons after 24 h treatment with OA 50 μM, OA+FAC, OA+N-acetylcysteine (NAC, 1 mM), or OA+FAC+NAC. **(F)** Primary neurons *Acc* expression after 24 h treatment under the conditions in **(E)**. Points denote biological classical replicates; bars show mean ± s.e.m. Two-group comparisons used two-tailed unpaired Student's t-test; multi-group comparisons used one-way ANOVA with Tukey's post hoc test. $*P \leq 0.05$, $**P \leq 0.01$, $***P \leq 0.001$. Scale bars, 100 μm.
